# Tau aggregates are RNA-protein assemblies that mis-localize multiple nuclear speckle components

**DOI:** 10.1101/2021.01.27.428450

**Authors:** Evan Lester, Felicia K. Ooi, Nadine Bakkar, Jacob Ayers, Amanda L. Woerman, Joshua Wheeler, Robert Bowser, George A. Carlson, Stanley B. Prusiner, Roy Parker

## Abstract

Tau aggregates contribute to neurodegenerative diseases including frontotemporal dementia and Alzheimer’s disease (AD). Although RNA promotes tau aggregation *in vitro*, whether tau aggregates in cells contain RNA is unknown. We demonstrate in cell culture and mouse brains that both cytosolic and nuclear tau aggregates contain RNA, with enrichment for snRNAs and snoRNAs. Nuclear tau aggregates colocalize with and alter the composition, dynamics, and organization of nuclear speckles, which are membraneless organelles involved in pre-mRNA splicing. Moreover, several nuclear speckle components, including SRRM2, mislocalize to cytosolic tau aggregates in cells, mouse brains, and patient brains with AD, frontotemporal dementia (FTD), and corticobasal degeneration (CBD). Consistent with these alterations we observe the presence of tau aggregates is sufficient to alter pre-mRNA splicing. This work identifies tau alteration of nuclear speckles as a feature of tau aggregation that may contribute to the pathology of tau aggregates.

## Introduction

Fibrillar aggregates of the microtubule associated protein tau (tau) are seen in numerous neurodegenerative diseases collectively referred to as tauopathies (Orr et al., 2017). Tauopathies have a variety of etiologies ranging from mutations in tau that promote its aggregation, such as in the inherited frontotemporal dementia with parkinsonism-17 (FTDP-17), to environmental triggers such as head trauma giving rise to chronic traumatic encephalopathy (CTE), to the incompletely understood link between beta-amyloid and tauopathy in Alzheimer’s disease (AD) (Aoyagi et al., 2019; Goedert et al., 1988; Wischik et al., 1988).

Several lines of evidence suggest that the formation and propagation of tau oligomers or aggregates is a key driver of toxicity in tauopathies. First, mutations that promote tau aggregation are causative in FTDP-17 (Goedert and Spillantini, 2000). Second, the rate of cognitive decline in AD is closely related to the rate of tau aggregate formation (Hanseeuw et al., 2019). Third, tau aggregates and tauopathy can be transmitted by inoculation in cells and mice (Aoyagi et al., 2019; Kaufman et al., 2016; Sanders et al., 2014; Woerman et al., 2016). Induction of tau aggregates in cell models also can be toxic (Sanders et al., 2014). Fourth, reduction of tau is neuroprotective in mouse models of AD (DeVos et al., 2018). Understanding how tau oligomers or aggregates form and how they induce neurotoxicity may lead to the development of therapeutics for numerous neurodegenerative diseases.

Tau is present in the human central nervous system as six splice isoforms. These isoforms differ in the number of N terminal inserts—0N, 1N, or 2N—and the number of microtubule repeat binding domains (RD)—3R or 4R (Buée et al., 2000; Park et al., 2016). The N terminal inserts have been shown to impact tau’s localization, interactions with membranes, spacing between microtubules, and signal transduction (Brandt et al., 1995; Chen et al., 1992; Lee et al., 1998; Liu and Götz, 2013). The positively charged RD has been shown to form the core of the amyloid fibrils present in the brains of patients with tauopathies and this is also where the majority of disease causing mutations are found (Buée et al., 2000; Falcon et al., 2018, 2019; Fitzpatrick et al., 2017; Goedert, 2005; Wegmann et al., 2013; Zhang et al., 2020).

Several observations suggest RNA may affect the formation of tau aggregates. First, tau binds RNA (Dinkel et al., 2015; Schröder et al., 1984; Wang et al., 2006; Zhang et al., 2017). Second, *in vitro* RNA promotes the conversion of soluble tau into insoluble aggregated tau, possibly because the negatively charged phosphate backbone of RNA can neutralize the positively charged RD of tau (Ambadipudi et al., 2017; Dinkel et al., 2015; Kampers et al., 1996). Third, tau immunopurifies with a number of RNA binding proteins in both the aggregated and unaggregated states (Bai et al., 2013; Broccolini et al., 2000; Gunawardana et al., 2015; Hales et al., 2014a, 2014b; Hsieh et al., 2019; Meier et al., 2016). Fourth, tau aggregates in AD and Pick’s disease have been found to stain positive for RNA using RNA dyes (Ginsberg et al., 1997, 1998). Finally, analysis of the RNAs interacting with tau in an unaggregated state by iCLIP suggests that tau preferentially interacts with tRNAs (Zhang et al., 2017). Thus, important questions are whether pathological tau aggregates contain RNA, and if so, what is the nature of those RNAs and what are the possible physiologic or pathologic consequences of their interaction?

Herein, we investigated the RNA composition of tau aggregates in both cell culture and mouse model systems. Similar to earlier results, we found that tau aggregates form in the cytosol and the nucleus (Bukar Maina et al., 2016; Gil et al., 2017; Rady et al., 1995; Sanders et al., 2014; Ulrich et al., 2018). We found that both cytosolic and nuclear tau aggregates contain RNA and are enriched for RNAs involved in RNA splicing and modification including snRNAs and snoRNAs, as well as repetitive Alu RNAs. We also found that nuclear tau aggregates contain snRNAs and are concentrated in, and alter the composition, organization and dynamics of, splicing speckles, which are non-membranous assemblies of RNA and protein containing nascent RNA transcripts and splicing machinery (Galganski et al., 2017). Surprisingly, we discovered that the serine arginine repetitive matrix protein 2 (SRRM2), a protein component of splicing speckles, mislocalizes from nuclear splicing speckles to cytosolic tau aggregates in cellular models of tauopathy, tauopathy mouse models, and patients with AD, frontotemporal lobar degeneration (FTLD), and corticobasal degeneration (CBD). These extensive interactions of tau with splicing speckles and the splicing machinery correlate with splicing alterations seen in cells that form tau aggregates. This is notably similar to how cytosolic sequestration of RNA binding proteins such as TDP-43 and FUS in amyotrophic lateral sclerosis (ALS) can lead to alterations in nuclear RNA processing promoting neurodegeneration (Lagier-Tourenne et al., 2012; Polymenidou et al., 2011).

## RESULTS

### Cytosolic and nuclear tau aggregates contain RNA

To determine whether tau aggregates contain RNA, we employed a previously developed HEK293 tau biosensor cell line (Holmes et al., 2014; Sanders et al., 2014). The HEK293 biosensor cells express the 4R repeat domain (RD) of tau with the P301S mutation tagged with either cyan-fluorescent protein (CFP) or yellow fluorescent protein (YFP). Fluorescent tau aggregates can be induced in these HEK293 cells via lipofection of preformed non-fluorescent tau aggregates isolated from the brains of mice expressing 0N4R tau with the P301S mutation (P301S mice, Tg2541) (Holmes et al., 2014; Sanders et al., 2014). As previously seen (Sanders et al., 2014), we observed fluorescent tau aggregates in both the cytosol and the nucleus of the HEK293 cells following transfection of clarified brain homogenate from mice expressing P301S human tau, but not from mice expressing wild-type (WT) tau (WT mice, Tg21221), which do not develop tauopathy (Fig. 1A-B). Nuclear tau aggregates are not an artifact of the truncated K18 tau expressed in HEK293 cells since we also observed the formation of both nuclear and cytosolic tau aggregates in a tau seeding model expressing full length P301S 0N4R tau-YFP in H4 neuroglioma cells (Fig. S1A, Supp. video 3). Consistent with these fluorescent bodies being insoluble tau aggregates, fluorescence recovery after photobleaching (FRAP) revealed that both nuclear and cytosolic tau aggregates are immobile and do not recover after photobleaching (Fig. S1B, C).

**Figure 1:**
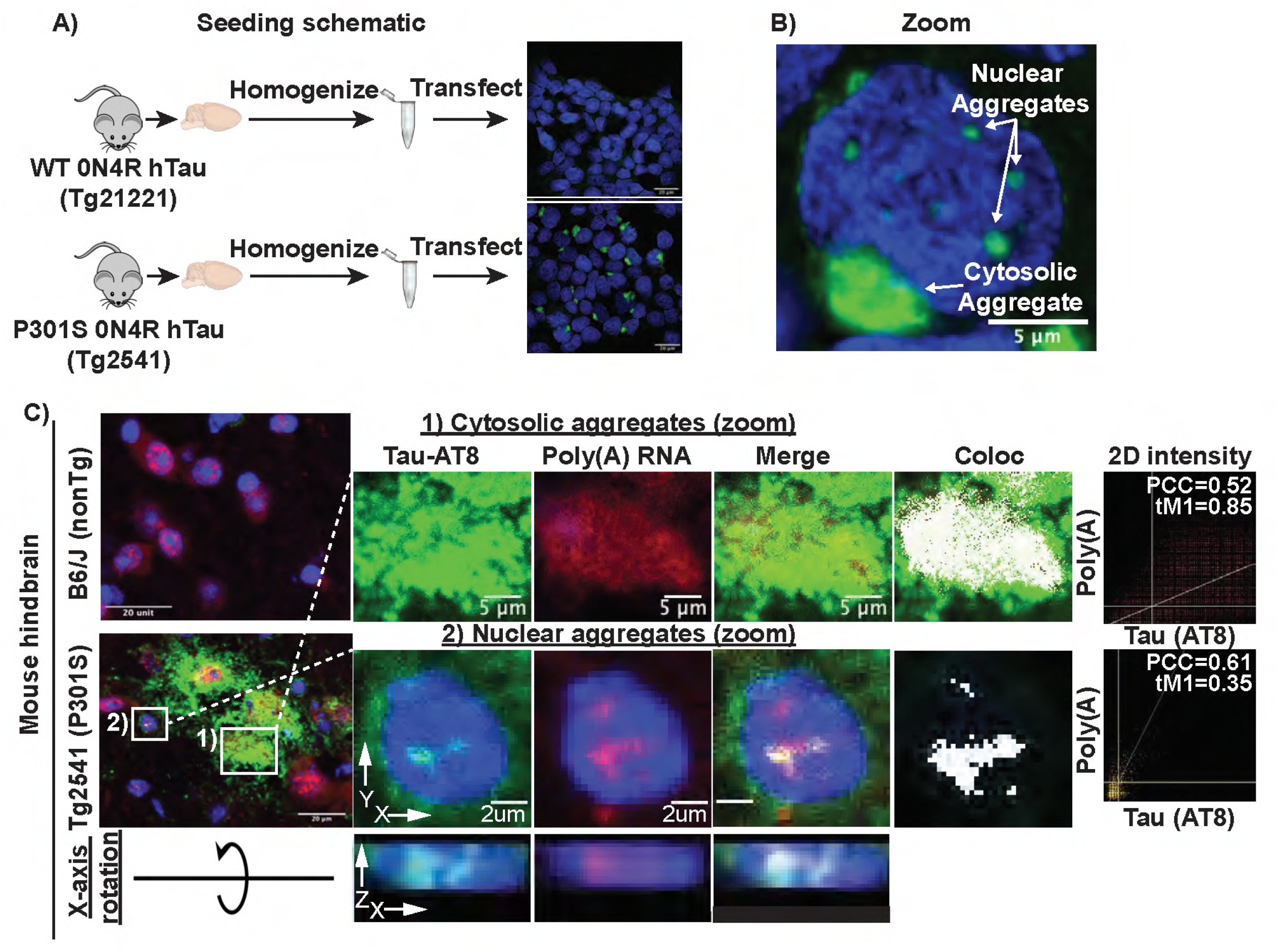
Tau biosensor cell schematic and tau aggregates in mice contain poly(A) RNA. Tau aggregates in HEK293 biosensor cells and mouse brain contain poly(A) RNA. **(A)** Schematic showing experimental design of tau seeding in HEK293 biosensor cells. Brain homogenate from mice expressing either WT (rTg21221) or P301S (rTg2541) 0N4R tau was homogenized, clarified by successive centrifugation and transfected into HEK293 cells expressing tau K18 (4R repeat domain) tagged with either CFP or YFP. Only cells transfected with P301S homogenate formed bright fluorescent aggregates. **(B)** Tau aggregates form in both the nucleus and the cytosol following transfection of P301S tau homogenate. **(C**) Cytosolic and nuclear tau aggregates contain poly(A) RNA in mouse brain. White pixels in Coloc image show pixels above the Costes determined thresholds in 2D intensity plots (PCC = Pearson correlation coefficient and tM1= thresholded manders colocalization (% of tau pixels above threshold that colocalize with poly(A) pixels above threshold)). X-axis rotation shows AT8 and oligo(dT) staining within the nucleus of mouse Tg2541 cells.

Using fluorescence in-situ hybridization (FISH) for poly(A) RNA we observed that cytosolic and nuclear tau aggregates showed 1.5- and 1.72-fold enrichment of poly(A)+ RNA staining, respectively (Figure S1D). We also examined the presence of poly(A) RNA in tau aggregates in the brains of 6 month old P301S mice, in which transmissible tau aggregates start to form at 1.5 months (Holmes et al., 2014; Yoshiyama et al., 2007). Unlike in humans, where tau pathology develops in the frontal cortex, tau pathology in P301S mice tau pathology develops primarily in the hindbrain (Johnson et al., 2017). Nuclear tau and tau aggregates have been previously observed in the brains of mice and humans, however their function and relevance to disease is poorly understood (Bukar Maina et al., 2016; Gil et al., 2017; Jiang et al., 2019; Liu and Götz, 2013; Montalbano et al., 2020; Rady et al., 1995; Ulrich et al., 2018). We observed nuclear tau aggregates in the hindbrain stain strongly for poly(A) RNA (Fig. 1C, S1E, S7A). We also observed a redistribution of poly(A) signal to overlap with the cytosolic tau aggregates in P301S mouse brains (Fig. 1C, S1E, S7A). Thus, in both mouse and cellular models of tau pathologies, tau forms cytosolic tangles and nuclear puncta that contain RNA.

### Tau aggregates in HEK293 cells and mouse brains are enriched for snRNAs and snoRNAs

To determine the identity of the RNAs present in tau aggregates, we first purified tau aggregates from HEK293 tau biosensor cells using differential centrifugation and fluorescent activated particle sorting (FAPS) and then sequenced the associated RNA (Fig. S2A-C). SYTO17 staining of the lysates post-sorting confirmed that RNA remained associated with the tau aggregates (Fig. S2A). By comparing the abundance of RNAs in total RNA and tau aggregates, we observed that tau aggregates contain a diverse transcriptome (Fig. 2A) and were enriched for small non-coding RNAs, particularly snoRNAs and minor snRNAs (Fig. 2B). Some mRNAs were also enriched in tau aggregates, notably mRNAs coding for voltage gated calcium channel complex, histone proteins, centrosomal proteins, and proteins involved in splicing regulation (Fig. S2D-E).

**Figure 2:**
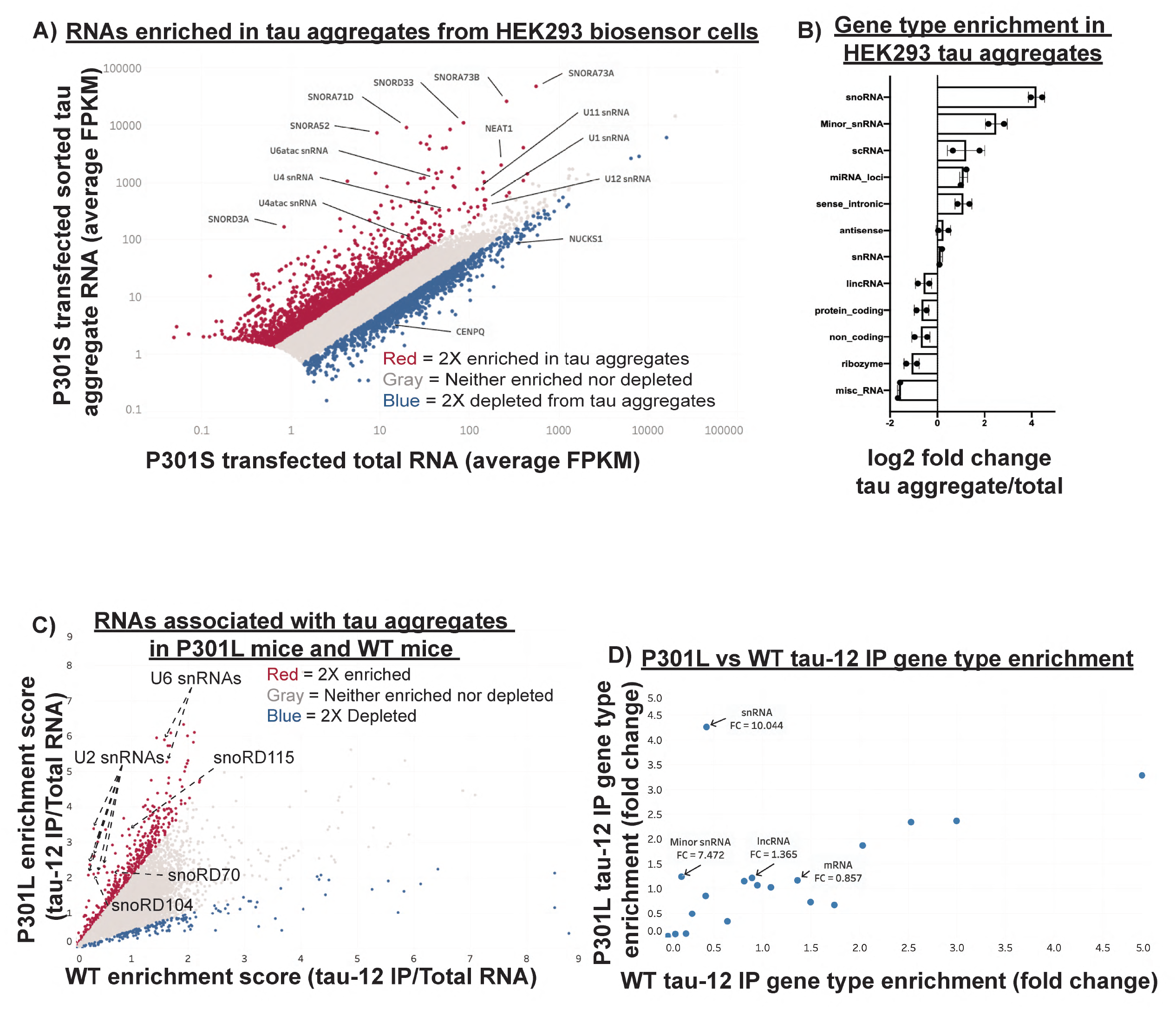
The RNA composition of tau aggregates in cellular and mouse tauopathy model systems. The RNA composition of tau aggregates in cellular and mouse tauopathy model systems. **(A)** Scatter plot of RNA sequencing showing average FPKM of two replicates for tau aggregate associated RNA and total RNA. Genes in red are two-fold enriched in tau aggregates and genes in blue are two-fold depleted from tau aggregates. Genes with fewer than 5 FPKM were removed from the analysis due to low coverage. **(B)** Fold change in the percentage of total FPKM for each gene type between the tau aggregate RNA and total RNA. Percentage of total FPKM was calculated by grouping genes using the Ensembl GRCh38.p13 biomart gene types. **(C)** Scatter plot of RNA sequencing data from mouse brain tau aggregate isolation in P301L and WT mice. Enrichment scores were calculated by dividing the insoluble tau IP FPKM by the total RNA FPKM for each replicate. Genes in red are two-fold enriched and genes in blue are two-fold depleted from the P301L sarkosyl insoluble tau aggregates. **(D)** Gene type enrichment in the P301L and WT samples. Fold change for each gene type was calculated by dividing the percentage of total FPKM made up by each gene type in the insoluble tau-IP by the percentage of total FPKM made up by each gene type in the total RNA. The numbers below the gene type names indicate the P301L/WT enrichment.

Analysis of RNAs expressed from multicopy genes using RepEnrich (Criscione et al., 2014) showed enrichment of RNAs from the multicopy snoRNAs (U3, U17, and U8) and the multicopy snRNAs (U2 and U1) (Figure S2F-G). Consistent with this observation, U1 snRNA has previously been observed to be enriched in AD tau aggregates by PCR (Hales et al., 2014a). Additionally, this analysis showed some enrichment of tRNAs, as previously observed with non-aggregated tau (Zhang et al., 2017). We also observed enrichment of RNAs from specific types of transposable elements, namely the hAT-Tip100 family of DNA transposable elements and Alu elements (Fig. S2F-G).

### Tau aggregates in P301L mouse brain are also enriched for snRNAs

To investigate whether tau aggregates in mouse brains contain similar RNAs to tau aggregates identified in HEK293 cells, we fractionated mouse brains with a 1% sarkosyl extraction followed by tau immunoprecipitation (IP) using the tau-12 antibody to isolate tau aggregates as previously described (Diner et al., 2017) (Fig. S2H). Western blot analysis showed that the sarkosyl extraction enriched for insoluble tau in the P301L mice (rTg4510) but not in the WT mice (rTg21221) (Fig. S2I). To identify RNAs specifically enriched in the aggregated tau fraction, we compared the enrichment of RNAs in the insoluble tau-12 IP relative to total RNAs between the WT and P301L mice (Fig. 2C).

Analysis of RNAs enriched in tau aggregates isolated from mouse brain relative to total RNA revealed an enrichment of specific RNAs, including snRNAs and snoRNAs in the P301L tau IP samples (Fig. 2C, D). This is similar to the RNA composition of tau aggregates isolated from HEK293 cells (Fig S2K). For example, we observed that snoRD115, snoRD104, snoRD70, U2 snRNA, and U6 snRNA, were enriched in the P301L insoluble tau fraction (Fig. 2C). In contrast to the HEK293 tau aggregates, only particular snoRNAs were enriched. This could be due to a variety of factors including differences in the isoforms of tau expressed in the two models (RD of tau in the HEK293 cells versus full-length 0N4R tau in the rTg4510 mice), the mutation in tau itself (P301S in the HEK293 biosensor cells and P301L in the rTg4510 mice), or differences in the RNA expression profiles of HEK293 cells and mouse neurons. The sarkosyl-insoluble fraction from both the P301L and WT mice revealed little to no enrichment of snRNAs or minor snRNAs suggesting that RNAs enriched in the tau-12 IP are interacting with aggregated tau rather than just enriched in the insoluble fraction (Fig. S2J). Similar to HEK293 cells, we observed some mRNAs enriched in tau aggregates from mouse brain (Figure S2L). Interestingly, some of the most enriched mRNAs are components of the centrosome (PCNT, Cep250, Cep164, Cep131) (Delaval and Doxsey, 2010; Graser et al., 2007), which is in agreement with previous work that has observed cytosolic tau aggregates concentrating at the centrosomes (Sanders et al., 2014; Santa-Maria et al., 2012). mRNAs coding for centrosomal proteins, such as PCNT, have previously been described to be present at the centrosome where they are locally translated and could become ensnared in tau aggregates (Sepulveda et al., 2018). Taken together, our results demonstrate that isolated pathological tau aggregates in mouse brains are enriched for similar types of RNAs as the tau aggregates in HEK293 cells including snRNAs and snoRNAs.

### Enriched RNAs localize to tau aggregates by fluorescence in-situ hybridization

We used FISH to examine if RNA enriched in tau aggregates identified by RNA sequencing localized to cytosolic and/or nuclear tau aggregates. We performed FISH for two enriched snRNAs (U2 and RNU6ATAC), two enriched snoRNAs (snoRA73B and snoRD3A), two depleted mRNAs (CENPQ and NUCKS1), and for the enriched Alu family of multicopy RNAs (Fig. 3, S3). Interestingly, we observed that enrichment of specific RNAs differed with respect to the localization of tau aggregates. Specifically, snoRD3A had a 2.05-fold enrichment into cytosolic tau aggregates relative to bulk cytosol, while nuclear tau aggregates had a 1.35-fold enrichment of snoRD3A relative to bulk nucleoplasm (Fig. 3A). For U2 snRNA, cytosolic and nuclear tau aggregates had roughly the same fold enrichment over their respective compartments (1.48 and 1.42) yet the absolute intensity of U2 snRNA in nuclear tau aggregates was 1.87-fold higher than that of the cytosolic tau aggregates (Fig. 3B). In cells without tau aggregates, U2 snRNA localizes into discrete nuclear foci called splicing speckles, which are non-membranous RNA-protein assemblies containing factors involved in mRNA splicing (Kota et al., 2008; Wagner et al., 2004; Zhang et al., 2016). Nuclear colocalization of the splicing speckle associated snRNAs with tau implies that nuclear tau aggregation is occurring in splicing speckles (see below). Consistent with this observation, we also see colocalization of the enriched RNU6ATAC snRNA in nuclear tau aggregates (Fig. S3A).

**Figure 3:**
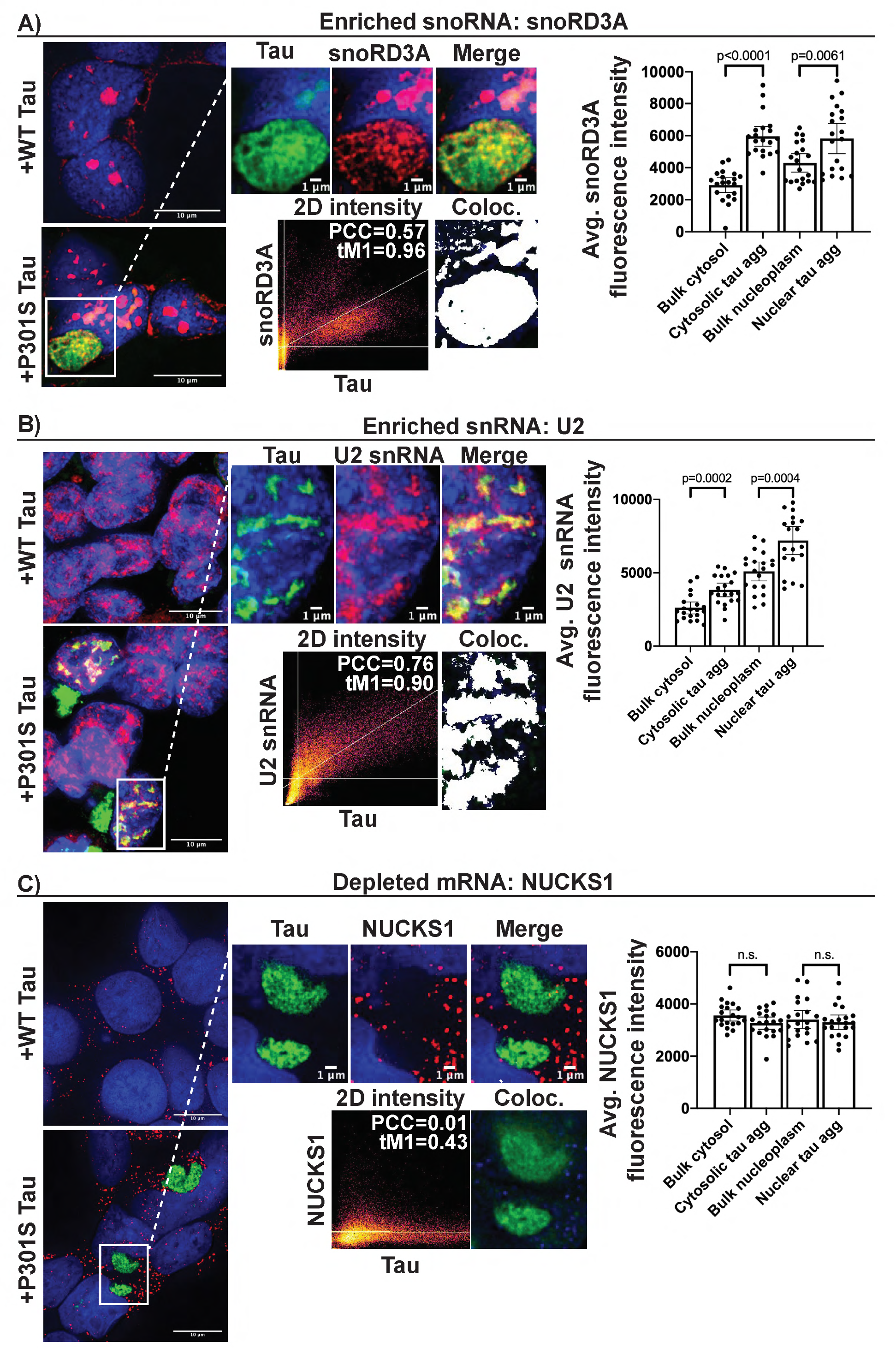
FISH for RNAs in HEK293 tau biosensor cells identified by sequencing. Line intensity plots and intensity quantification show enrichment of snoRD3A **(A)** and U2 snRNA **(B)** in both the nucleus and cytosol. We observed no enrichment of the depleted mRNA, NUCKS1 **(C)**, into nuclear or cytosolic tau aggregates. Bar graphs show quantification of FISH fluorescence intensity within nuclear and cytosolic tau aggregates in relation to bulk cytosol and nucleoplasm (n = 20 aggregates). Significance was determined using an unpaired two-tailed t-test. White pixels in Coloc image show pixels above the Costes determined thresholds in 2D intensity plots (PCC = Pearson correlation coefficient and tM1= thresholded manders colocalization (% of tau pixels above threshold that colocalize with red pixels above threshold)).

We also performed FISH for an Alu consensus sequence present in the Alu RNAs enriched from the sequencing data (Fig. S2F-G) and found that Alu signal enriched into both nuclear and cytosolic tau aggregates (Fig. S3C). Alu enrichment was greater in cytosolic aggregates (1.56-fold; p<0.0001) than in nuclear aggregates (1.23-fold; p=0.02). In the nuclear aggregates, Alu intensity was greatest on the periphery of the aggregates. The depleted mRNA NUCKS1 did not show significant intensity enrichment into either nuclear or cytosolic tau aggregates (Fig. 3C). Similarly, no positive localization correlation could be found for the depleted mRNA CENPQ and tau (Fig. S3D). Thus, tau aggregates contain, and are enriched for, a diverse set of specific RNAs.

### Nuclear tau aggregates colocalize with splicing speckles in HEK293 cells

Due to the bias in both the HEK293 and mouse tau aggregate transcriptomes towards nuclear snRNAs and snoRNAs, we explored whether nuclear tau aggregates localized to the nucleolus, where snoRNAs are concentrated, or splicing speckles, which are enriched in snRNAs (Spector and Lamond, 2011).

Three lines of evidence suggest that nuclear tau aggregates colocalize with splicing speckles. First, the canonical splicing speckle antibody SC35 localized with nuclear tau aggregates by immunofluorescence (IF). In IF, the SC35 antibody recognizes the splicing speckle protein, SRRM2 (Ilik et al., 2020), and in agreement with this specificity, an SRRM2-halo fusion protein also localizes to nuclear tau aggregates (Fig. 4A, Fig. S4A). Similarly, other components of nuclear speckles co-localize with nuclear tau aggregates (see below). Second, nuclear tau aggregates are enriched for poly(A) RNA (Fig. 1C, S1D-E, Supp. Video 1, 2) and splicing speckle associated RNAs including U2 snRNA (Fig. 3B, S2F) (Huang et al., 1994). Third, consistent with tau accumulating in splicing speckles and not with other nuclear RNA foci, nuclear tau aggregates do not colocalize with the nucleolar markers fibrillarin and RPF1 (Kiss, 2002) (Fig. S4B-C). In the brains of P301S (Tg2541) mice, we also observe pTau(S422) signal colocalizing with both SRRM2 and poly(A) in the nucleus (Fig. S7A). Thus, nuclear tau aggregates localize to SRRM2-positive nuclear splicing speckles in cell and mouse models of tauopathy.

**Figure 4:**
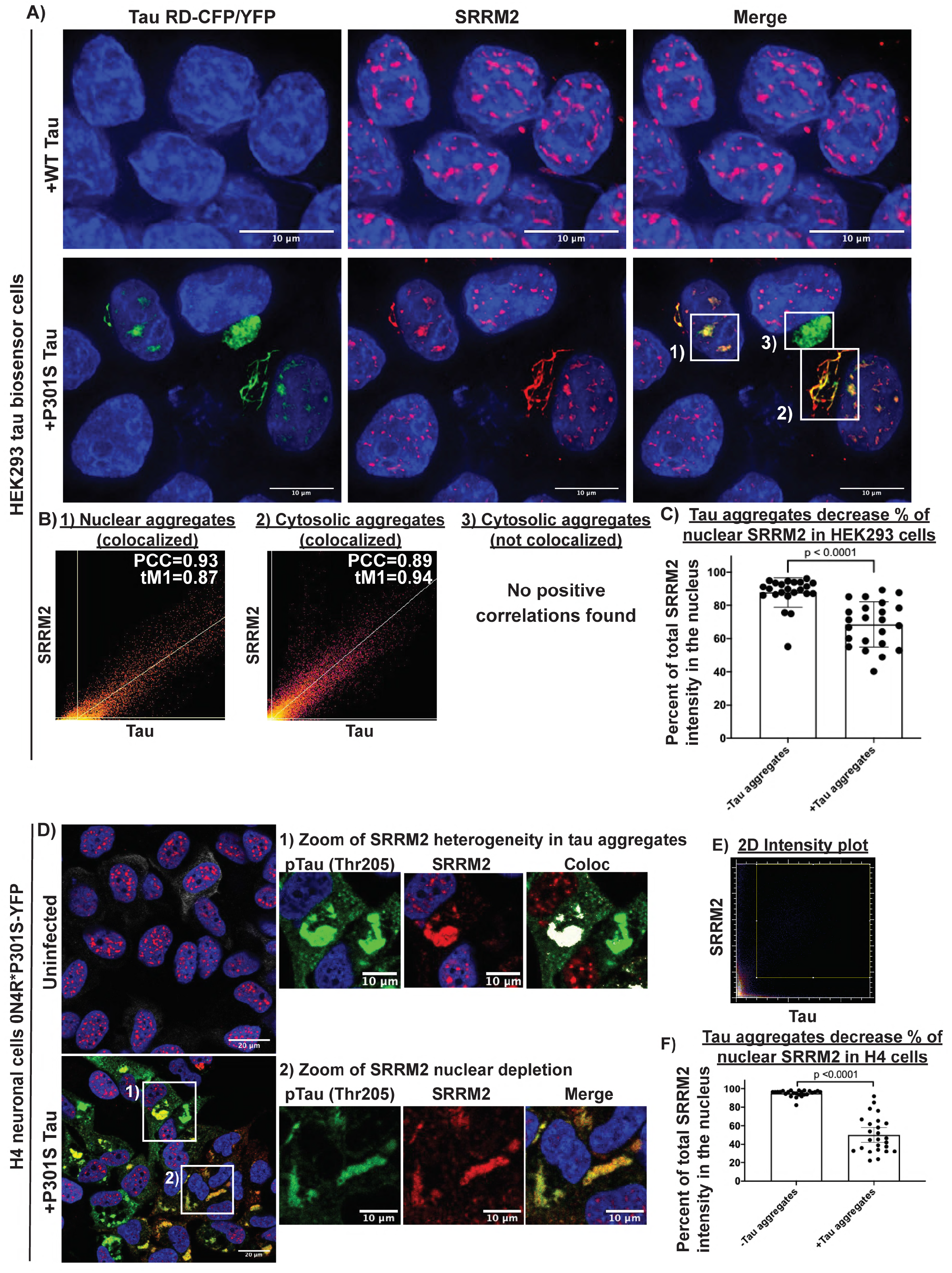
Tau aggregates colocalize with splicing speckles and mislocalize SRRM2 in tauopathy model systems. **(A)** Nuclear tau aggregates in HEK293 cells colocalize with SRRM2 (SC-35 antibody), a marker of splicing speckles. **(B)** Colocalization analysis showing the relationship between various tau aggregates and SRRM2 **1)** nuclear tau aggregates and splicing speckles marked by SRRM2, **2)** a cytosolic tau aggregates that colocalize with cytosolic SRRM2, and **3)** a cytosolic tau aggregate that does not colocalize with SRRM2. **(C)** Quantification of the percent of total SRRM2 intensity in the nucleus in HEK293 cells with and without tau aggregates (n = 23 cells). **(D)** Immunofluorescence of phospho-tau (Thr205) and SRRM2 in H4 neuronal cells expressing 0N4R*P301S-YFP tau +/- tau aggregates shows SRRM2 recruitment to tau aggregates is not dependent on phosphorylation at Thr205. **(E)** 2D intensity plot for the zoomed images showing two Thr205 positive tau aggregates, one that colocalizes with SRRM2 and one that does not colocalize with SRRM2. White pixels in Coloc image show pixels above the Costes determined thresholds in 2D intensity plot. **(F)** Quantification of percentage of SRRM2 in the nucleus in cells with and without tau aggregates (n = 25 cells). Images were quantified using CellProfiler and P-values were calculated using an unpaired two tailed t-test.

### Multiple nuclear speckle components re-localize to cytoplasmic Tau aggregates

While examining the co-localization of tau and SRRM2 in nuclear speckles in HEK293 cells, we observed that 77% of cytosolic tau aggregates also contained SRRM2 that relocalized to the cytosol (Fig 4A & S4D). Similar results were observed with an SRRM2-halo fusion protein (Fig. S4A). Quantification of SRRM2 IF in cytosolic and nuclear tau aggregates revealed an 11.12-fold and 2.10-fold enrichment over bulk cytosol and nucleoplasm respectively (Fig. S4E). Both the colocalization of nuclear tau aggregates in splicing speckles and the relocalization of SRRM2 to cytosolic tau aggregates were independent of the lipofectamine used to transfect tau into the cell models (Fig. S4F).

The accumulation of SRRM2 into cytosolic Tau aggregates was sufficient to deplete nuclear SRRM2 in HEK293, and an H4 neuroglioma cell line expressing a full length 0N4R P301S tau-YFP that forms fewer nuclear tau aggregates (Fig 4C, 4F). Moreover, we observed SRRM2 relocalized into cytoplasmic phospho-tau aggregates in cells without nuclear tau aggregates demonstrating the cytosolic accumulation is independent of nuclear tau aggregates (Fig. 4D). Interestingly, in H4 cells that accumulated SRRM2 in cytosolic tau aggregates, nuclear speckles formed with no change in the intensity of the SON protein, a nuclear speckle protein that does not accumulate in cytosolic tau aggregates (Fig. S4G-H). These observations argue that cytosolic tau aggregates deplete the nucleus of SRRM2, but do not prevent the formation of nuclear speckles.

To determine if other nuclear speckle proteins also re-localized into cytosolic tau aggregates to some extent, we utilized IF to measure the average intensity enrichment in nuclear and cytosolic tau aggregates. We observed that the SRRM2 paralogs, SRRM1 and SRRM3, did not accumulate in tau aggregates (Fig. 5A, S5A) indicating this accumulation is not shared between SRRM family members. Similarly, proteins with SR domains did not accumulate in cytosolic tau aggregates (SRSF1, SRSF2, and SRSF3) indicating that an SR domain is not sufficient for accumulation in cytoplasmic tau aggregates (Fig. 5A, S5A). Some, but not all, speckle components/splicing factors showed cytosolic tau co-location with PNN (a known binding partner of SRRM2), SFPQ, MSUT2, DDX39B, and DYRK1A showing the strongest enrichment scores (Figure 5A, S5A) (Zimowska et al., 2003). Thus, multiple nuclear speckle proteins involved in pre-mRNA splicing mis-localize to cytosolic tau aggregates with SRRM2, PNN, and SFPQ being the most strongly re-localized.

**Figure 5:**
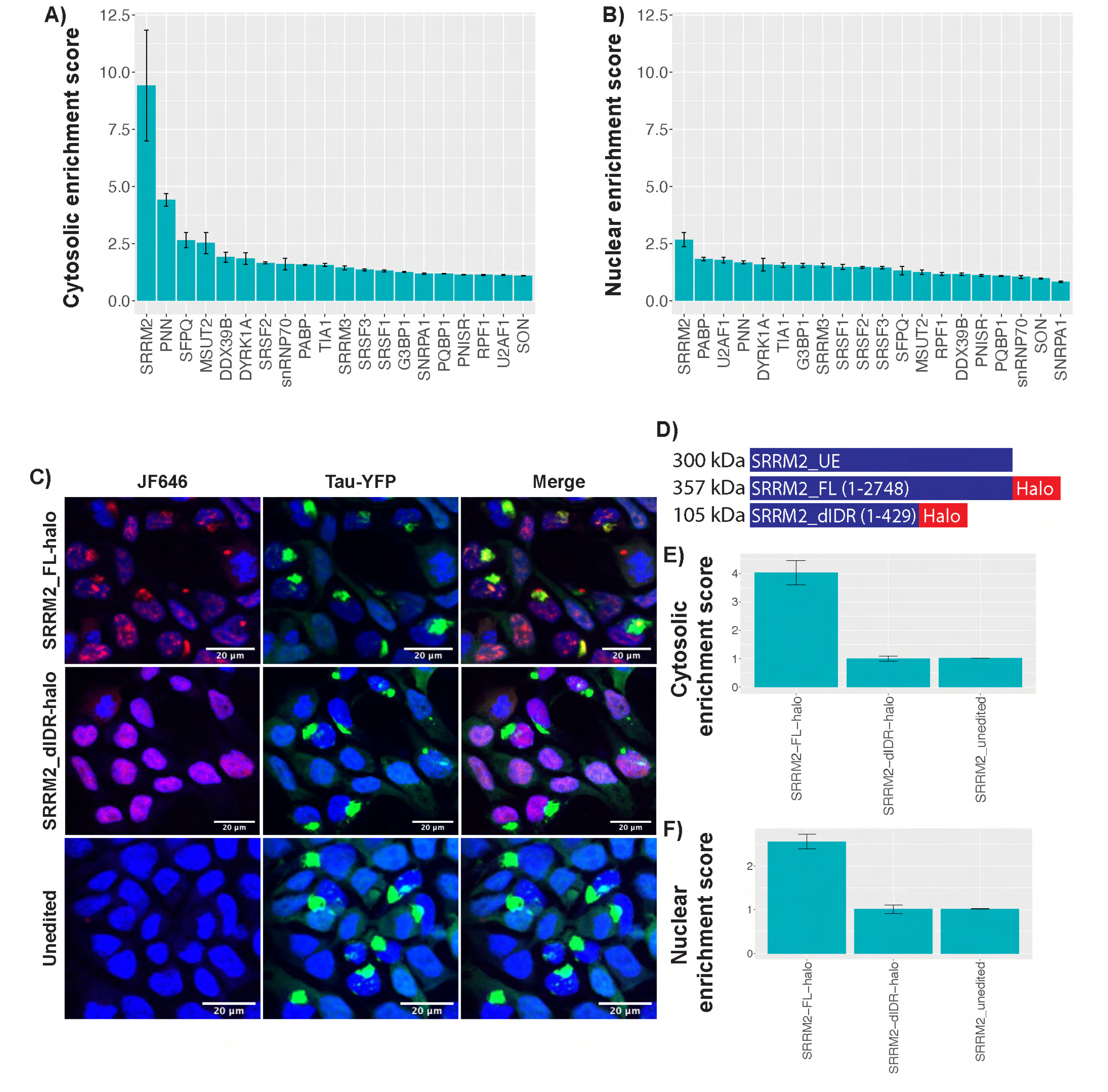
Other proteins that localize to tau aggregates and the C-terminal region of SRRM2 is responsible for localization to tau aggregates. **(A, B)** Cytosolic and Nuclear tau aggregates enrichment scores (Median intensity within tau aggregate/median intensity within cytosol or nucleus) for 20 proteins. Ilastik was used to segment images into the following categories: tau aggregates, nucleus, cytosol, and background. Segmentation masks were fed into CellProfiler to quantify the median intensity of the interrogated protein. N=25 images per condition, scale bar represents 95% confidence interval. **(C)** Images of cells CRISPR edited to express endogenous FL or dIDR SRRM2 tagged with halo. **(D**) schematic of SRRM2 constructs. **(E, F)** Cytosolic and nuclear enrichment scores for FL-Halo, dIDR-Halo, and unedited SRRM2.

Since neither SR domains nor the N-terminal conserved features of SRRM proteins appeared sufficient to recruit proteins to tau aggregates, we hypothesized that the C-terminal domain of SRRM2, which is comprised of an intrinsically disordered region (IDR) (Fig. S5B), might be responsible for SRRM2 recruitment to tau aggregates. This would be consistent with the trend that intrinsically disordered regions of proteins can promote their recruitment to membraneless organelles. To test this idea, we used the CRISPaint system to create two HEK293 tau biosensor cell lines that contained a halo tag inserted into endogenous SRRM2 (Ilik et al., 2020; Schmid-Burgk et al., 2016). These two cell lines were 1) a full length SRRM2 cell line referred to as SRRM2_FL-halo (insert at aa 2708), and 2) a cell line lacking the C-terminal IDR of SRRM2 referred to as SRRM2_dIDR-halo (insert at aa 430) (Fig. 5C, S5B-D). We induced tau aggregation in these cells and compared the average SRRM2 halo intensity within nuclear and cytosolic tau aggregates relative to the average intensity in the bulk nucleus or cytosol respectively.

We observed that SRRM2_FL-halo was recruited to tau aggregates, but that the SRRM2_ dIDR-halo was not (Figures 5C-E). This demonstrated that SRRM2 is recruited to tau aggregates by the disordered C-terminal domain rather than the structured N-terminal domain. Since the N-terminal domain of SRRM2 is sufficient for RNA binding and interactions with the core of the spliceosome (Grainger et al., 2009; Zhang et al., 2018), this result argues that SRRM2 is not recruited to cytosolic tau aggregates by binding RNA nor the core of the spliceosome.

### Tau aggregates alter the properties of nuclear speckles including pre-mRNA splicing

Since cytoplasmic tau aggregates depleted some nuclear speckle components, and tau aggregates can also form in nuclear speckles, we hypothesized that tau aggregate formation might alter the properties and function of nuclear speckles, which we examined in three experiments. First, given that nuclear speckles are highly dynamic structures (Rino et al., 2007) yet the tau aggregating in speckles was essentially static (Fig S1C), we examined if speckles with tau aggregates showed altered dynamics by performing FRAP on two tagged components of speckles, SRRM2 (Halo) and SRSF2 (mCherry). We observed that in the presence of tau aggregates, both speckle components showed an increase in the static component, and a reduced rate of recovery from FRAP (Fig 6A-B). This demonstrates that the presence of Tau aggregates in nuclear speckles changes their dynamics.

**Figure 6:**
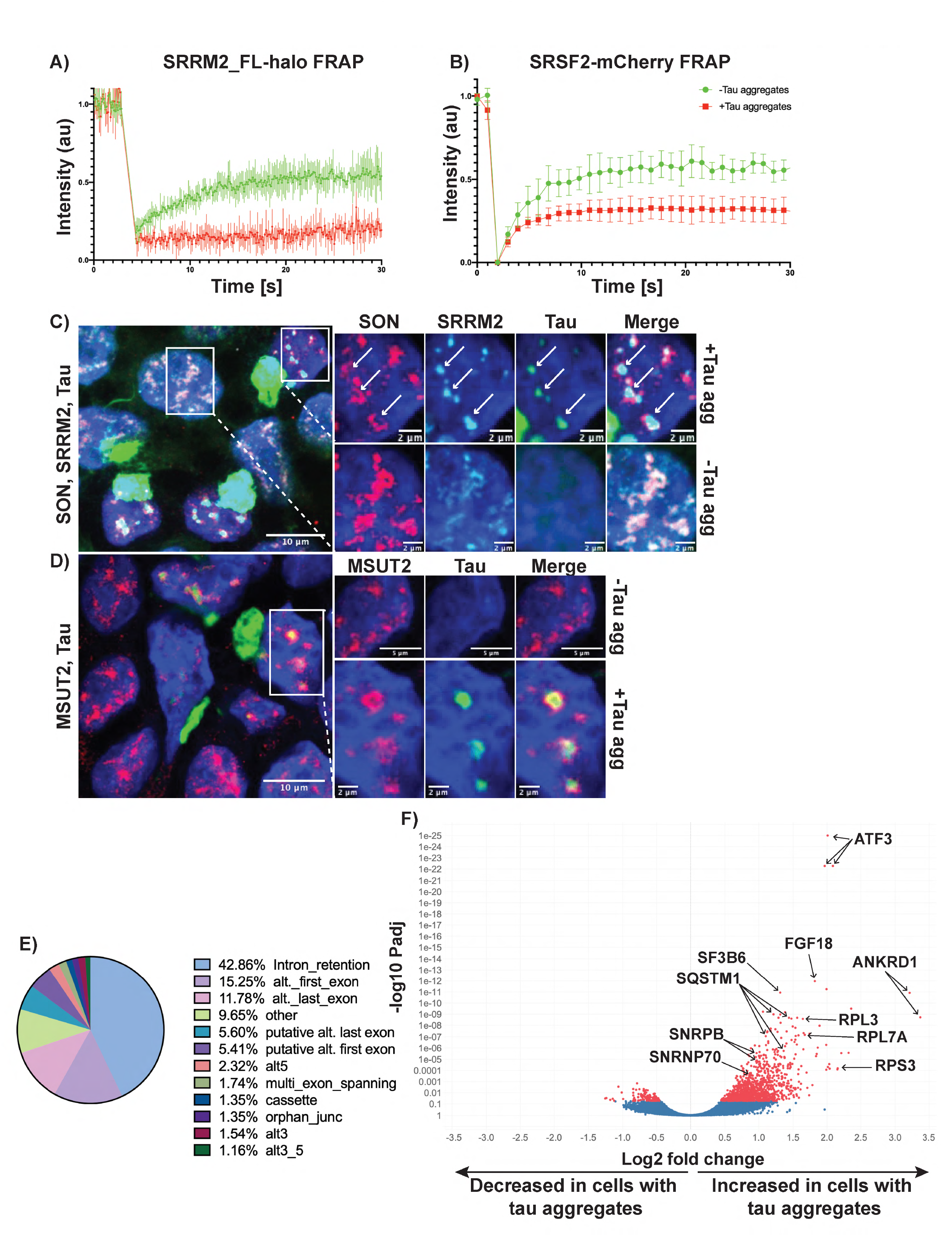
Tau aggregation alters dynamics and organization of splicing speckles and RNA splicing. **(A)** FRAP of SRRM2_FL-halo and **(B)** SRSF2-mCherry splicing speckles with and without tau aggregates (n=5). 405 nm laser was used to photobleach and fluorescence intensity was measured using the 647 nm channel for halo-JF647 and 561 nm for mCherry. **(C, D)** Images of P301S tau transfected cells showing nuclear speckle reorganization in the presence of aggregates. SON and MSUT2 move to the periphery of speckles, while SRRM2 remains in the center of speckles with the tau aggregate. **(E)** MAJIQ Analysis of splicing changes in cells with and without tau aggregates showing that most splicing changes are intron retention events. **(F)** Volcano plot showing differential intron retention in cells with tau aggregates quantified by iREAD and DEseq2 (Red = Padj < 0.05). Multiple points per gene are due to the multiple retained introns in those genes.

Second, the formation of tau aggregates in speckles alters the organization of speckle components. IF for SRRM2 and SON [a speckle protein that does not relocalize to cytosolic tau aggregates (Fig. 5A)] showed that the two proteins colocalize in speckles in the absence of tau aggregates. However, in the presence of nuclear tau aggregates, SRRM2 and tau colocalize in the center of speckles, while SON moves to the periphery and forms a ring-like structure around the aggregate (Fig. 6C). MSUT2, another protein that shows little relocalization to cytosolic tau aggregates, displayed a similar redistribution from the center of speckles to the periphery in the presence of tau aggregates. Interestingly, knockdown of MSUT2 has been shown to suppress tau toxicity in several model systems (Guthrie et al., 2011; Wheeler et al., 2019), suggesting that tau aggregates disrupt the spatial organization of speckles.

Third, since nuclear speckles are thought to modulate pre-mRNA splicing (Spector and Lamond, 2011), we performed RNA-Seq on the same HEK293 cells with and without tau aggregates to determine if the presence of tau aggregates could alter splicing. We then investigated splicing patterns using two analyses: MAJIQ and iREADs (Li et al., 2020; Vaquero-Garcia et al., 2016). Specifically, we observed using MAJIQ, at a ΔPSI threshold of 0.1 and confidence threshold of 0.95, we identified 305 local splicing variations in 226 genes that are differentially spliced (Supplemental Table 4). Examination of the types of local splicing variations revealed that the largest categories were intron retention (42.86%), alternative first exons (15.25%), and alternative last exons (11.78%) (Fig 6E MAJIQ splicing diagrams and IGV raw read counts for one example, ATF3, are provided in Fig. S6A-B). Due to the abundance of intron retention events, we used iREAD (intron REtention Analysis and Detector) to better quantify differential intron retention between cells with and without tau aggregates. Reads that fully or partially overlap annotated introns were then used for differential expression analysis using DEseq2 (Li et al., 2020) (Love et al., 2014). We found that at a Padj < 0.05 there were 1,225 introns in 641 genes that were retained in cells with tau aggregates and 120 introns in 86 genes that were retained in cells without tau aggregates (Fig. 6F). Pre-mRNAs with retained introns in cells with tau aggregates cluster in genes affecting apoptosis and splicing associated protein (ASAP) complex, ribosome, and RNA splicing and processing (Fig. S6C). Thus, the formation of tau aggregates in cells is sufficient to induce changes in pre-mRNA splicing and is expected to have significant biological impact. Since patients with tauopathies and model systems with tau mutations show changes in pre-mRNA splicing patterns in the brain (Apicco et al., 2019; Hsieh et al., 2019; Raj et al., 2018), an alteration in splicing due to tau aggregate formation may contribute to these splicing changes (see discussion).

### SRRM2 is depleted from the nucleus and relocalized to cytosolic tau neurofibrillary tangles in mouse and human tauopathies

Since SRRM2 was the most highly enriched protein identified, we examined whether cytosolic tau aggregates also contain SRRM2 in mice via IF, using the SC35 antibody on brains from WT B6/J mice, or the Tg2541 mice. In B6 control mice, SRRM2 predominantly localized to poly(A)+ nuclear splicing speckles (Fig. S7A). In contrast, we observe SRRM2 re-localized from the nucleus into cytosolic phospho-tau aggregates in the P301S-expressing Tg2541 mice (Fig. S7A). Consistent with these observations, phosphorylated SRRM2 has been previously observed to be relocalized to the cytosol in 5X FAD mouse brains, however the association of SRRM2 with tau aggregates was not reported (Tanaka et al., 2018). Thus, SRRM2, and potentially other speckle components, relocalize and sequestered into cytosolic tau aggregates in both cell culture models and in tauopathy mice.

To examine if SRRM2 is mis-localized in human tauopathies, we performed IF on tauopathy patient brains. We observed that in patients with the primary tauopathy CBD, SRRM2 was present in tau-containing cytosolic aggregates in the form of neuropil threads, whereas SRRM2 localized to nuclear splicing speckles in aged-matched heathy control patients (Figure 7A, patient demographics in Supplemental Table 3). Quantification of CBD and age matched control images revealed that the average nuclear SRRM2 signal was significantly lower in the CBD brains compared to the controls (Fig 7B). We also observed that SRRM2 was re-localized from the nucleus into the cytosol in the frontal cortex of multiple AD and FTLD patient brains (n=4 AD and n=4 FTLD), but not age-matched control brains (n=4, Figure 7C, S7B, patient demographics in Supp. Table 3). These results show that cytosolic SRRM2 marked by the SC35 antibody is a histopathological feature seen across three distinct human tauopathies.

**Figure 7:**
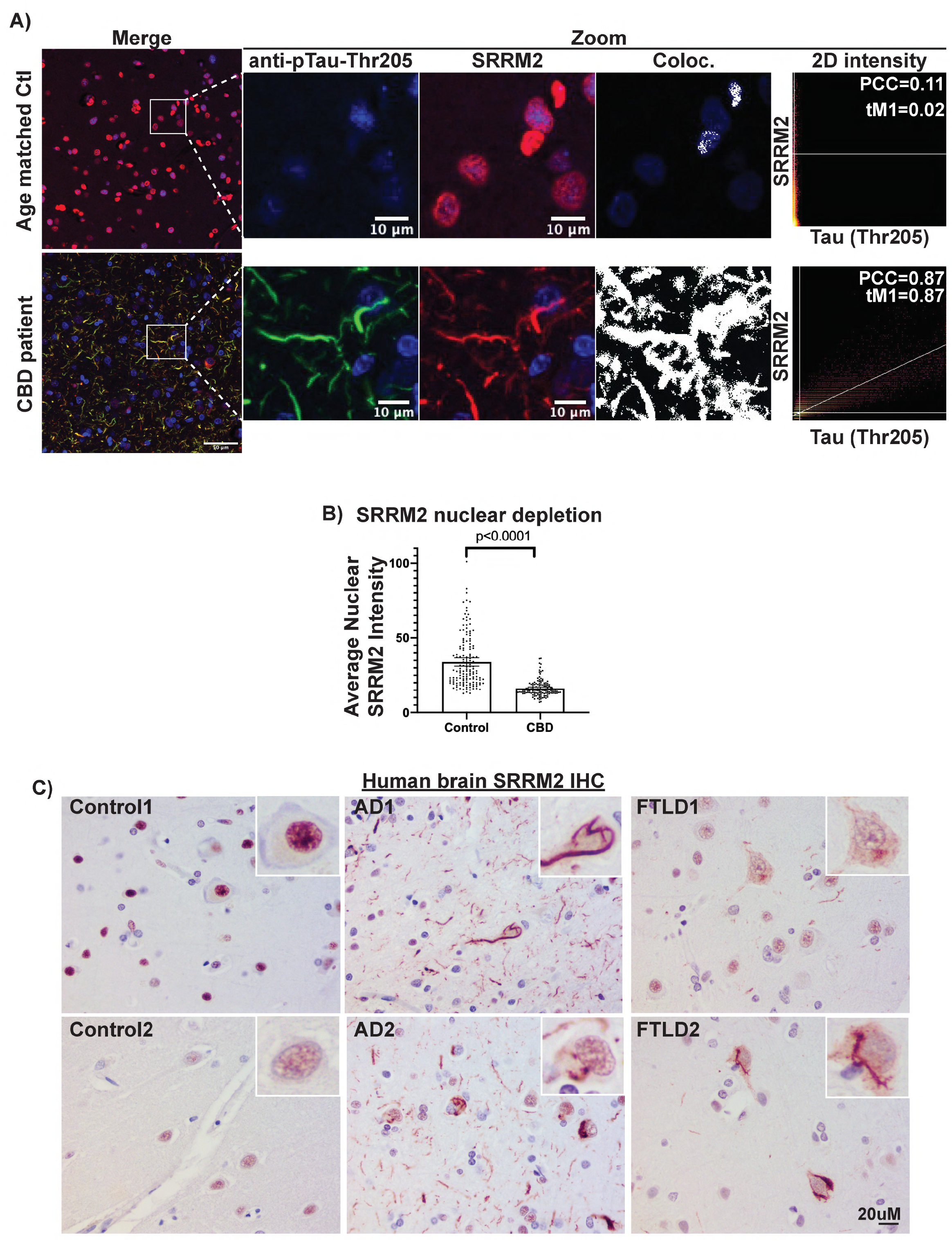
SRRM2 is relocalized to cytosolic tau aggregates in human tauopathies. **(A)** IF and colocalization analysis showing hyperphosphorylated tau (pTau-Thr205) colocalizes with SRRM2 (SC35) in the cytosol of CBD patient brain while SRRM2 is localized to the nucleus in age matched control brain. **(B)** Quantification of the average nuclear SRRM2 intensity showing a significant decrease in the setting of CBD relative to age matched control. Error bars are 95% confidence interval and p-values were calculated using an unpaired two tailed t-test. Immunohistochemisty of SRRM2 in human brains showing SRRM2 redistribution to the cytosol in AD and FTD brains but retains nuclear localization in control brains. Patient demographics and additional examples can be found in Fig. S5D Table S3.

## DISCUSSION

We present several lines of evidence that both cytosolic and nuclear tau aggregates contain RNA. First, in mice and HEK293 cells, nuclear and cytosolic tau aggregates both stained positive for poly(A) RNAs, indicating the presence of mRNAs or non-coding RNAs with poly(A) tails (Fig. 1, S1D-E, S7A). Second, purification and sequencing of tau aggregates from mouse brains or HEK293 cells demonstrated the presence and enrichment of specific RNAs, most notably snRNAs and snoRNAs (Fig. 2). Third, FISH for specific RNAs in HEK293 cells validated that our sequencing identified RNAs enriched in tau aggregates (Fig. 3). Although we have not yet examined the specific RNAs present in tau aggregates in human pathologies, tau aggregates in patient brains stain positive with acridine orange, a dye with specificity for RNA (Ginsberg et al., 1997, 1998). Based on these observations, we suggest that tau aggregates generally contain RNA, and the presence of specific RNA species may alter their formation and stability. The presence of RNA in tau aggregates may explain why tau aggregates and other RNA binding proteins can co-immunoprecipitate and/or co-localize (Apicco et al., 2018; Gunawardana et al., 2015; Hales et al., 2014a, 2014b; Maziuk et al., 2018; Meier et al., 2016; Vanderweyde et al., 2016).

We observed that tau aggregates are enriched for a number of different RNA species. Most notably, we observed enrichment of specific snRNA and snoRNAs in both HEK293 reporter cells and in mouse brains, although the specific snRNA/snoRNA species can vary between model systems (Fig. 2). We also observed the enrichment of repetitive RNAs (such as tRNAs, Alu elements, and satellite RNAs) and some mRNAs coding for proteins in the centrosome and spliceosome (Fig. S2D-L). As our library preparation protocol was not specifically designed to capture tRNAs, miRNAs, or rRNAs, our analyses may underestimate the abundance of these species (Motameny et al., 2010; Xu et al., 2019). The mechanisms by which specific RNAs are enriched in tau aggregates remains to be determined but could be due to tau’s intrinsic RNA binding specificity, the structure of the tau conformers, and/or the presence of specific RNAs at sites of tau aggregation such as snRNAs in nuclear speckles or mRNAs at the centrosome (Sepulveda et al., 2018). We suggest tau aggregates in cells could be considered a representative of the growing class of RNA and protein assemblies.

We provide evidence that nuclear tau aggregates form in splicing speckles and alter their properties, composition, and function. One critical observation is that nuclear tau assemblies are observed both in HEK293 cells and P301L mice that overlap with both protein and RNA markers of nuclear speckles (Fig. 4, S4, S7). Moreover, nuclear speckles that contain tau aggregates show altered dynamics of both SRRM2 and SRSF2 (Fig. 6A) demonstrating tau aggregation has altered their material properties. Moreover, tau accumulation in speckles also changes their organization with proteins partitioning into novel sub-domains of the assembly (Fig. 6C-D). Finally, since cytoplasmic tau aggregates accumulate multiple components of nuclear speckles, leading to their depletion from the nucleus, the presence of tau aggregates within cells alters the composition of nuclear speckles.

A critical question is the biological significance of the nuclear tau aggregates. In this light it is important to note we observed tau aggregates in both cell and mouse models of tauopathies demonstrating nuclear tau aggregates are not an artifact of cell line models (Fig. 1C, S1E, S7A). Moreover, we observe tau aggregates in nuclear speckles in both HEK293 cells expressing just the 4 repeat regions of tau, and in H4 neuroglioma cells expressing a full-length tau isoform (Fig. S1A) demonstrating tau accumulating in speckles is not an artifact of expressing truncated tau. Although tau is predominantly thought to be a cytosolic microtubule associated protein, numerous labs have observed aggregated and unaggregated tau in the nucleus of neuronal and non-neuronal cells (Bukar Maina et al., 2016; Gärtner et al., 1998; Gil et al., 2017; Jiang et al., 2019; Liu and Götz, 2013; Maj et al., 2010; Montalbano et al., 2020; Rady et al., 1995; Siano et al., 2019; Ulrich et al., 2018; Violet et al., 2014, 2015). Indeed, previous studies in HEK293 cells found that transfection of tau conformers that produced nuclear tau aggregates with a “speckled” phenotype were associated with greater cellular toxicity relative to those that only produced cytosolic tau aggregates (Sanders et al., 2014). In agreement, the suppressor of tau toxicity, MSUT2 (Guthrie et al., 2011; Wheeler et al., 2019), also localized to nuclear splicing speckles (Fig. 6D, S5A). Thus, tau’s interaction with splicing speckle components could be integral to the toxicity of tau aggregation.

A striking feature of our results is that cytoplasmic tau aggregates accumulate and mis-localize several proteins that normally accumulate in nuclear speckles (Fig. 4-6, S4-S7). The most striking of these is the SRRM2 protein, which can show an order of magnitude enrichment in cytosolic tau aggregates as compared to the bulk cytosol (Fig. 5A, S4E). Moreover, we also observed accumulation of SRRM2 in tau aggregates in mice, and in the human tauopathy, CBD (Figure 7A). Strikingly, we observed cytosolic mis-localization and/or nuclear depletion of SRRM2 in multiple tauopathies, including AD and FTLD (Fig. 7C, S7B). The consistent mis-localization of SRRM2 in cell line and animal models of tauopathy, as well as in deceased patient brain samples, argues the mis-localization of nuclear speckle proteins into cytosolic tau aggregates is a fundamental and consistent consequence of tau aggregation. An important issue in future work is to determine the mechanism of nuclear speckle component mis-localization and its biological consequences.

One likely consequence of altering the composition and dynamics of nuclear speckles would be to alter pre-mRNA splicing. It is well documented that pre-mRNA splicing is altered in tauopathy patient brains, including AD patients (Hsieh et al., 2019; Raj et al., 2018). Consistent with this finding, we demonstrate that the formation of tau aggregates in HEK293 cells is sufficient to induce alterations in alternative splicing and increase the number of significantly retained introns (Fig. 6E, S6). Thus, tau aggregates are sufficient to alter pre-mRNA splicing, although it should be noted that splicing alterations seen in disease tissue may be complicated by additional factors including multiple cell types and neuroinflammatory responses. Interestingly, many of these retained introns triggered by tau aggregates are in RNAs that code for proteins involved in RNA processing and ribosome biogenesis (Fig. S6A). Disruptions to these processes could lead to a pathologic cascade, potentially explaining the complex alterations in ribosome function and RNA processing that have been observed in AD patients (Hsieh et al., 2019; Q et al., 2005; Raj et al., 2018). These results raise the possibility that tau aggregation per se is responsible for some of the splicing changes seen in disease tissue.

The coaggregation of RNA and proteins in tauopathies is reminiscent of pathologic RNA-protein aggregates seen in other neurodegenerative and neuromuscular diseases, such as amyotrophic lateral sclerosis and inclusion body myopathy (Ramaswami et al., 2013; Taylor et al., 2016). Thus, the sequestration of RNAs and RNA binding proteins into pathologic aggregates may represent a shared pathophysiological feature across multiple degenerative diseases affecting diverse tissue types with a common feature being depletion of critical RNA processing factors from the nucleus leading to changes in RNA processing and gene expression.

## SUPPLEMENTAL FIGURE LEGENDS

**Supplemental Movies 1, 2, 3:** 3D rotation of a HEK293 cell nucleus expressing K18-YFP tau **(1)** and a mouse brain nucleus **(2)** showing intranuclear tau aggregates (green) (YFP tagged tau in HEK293 cells, and AT8 IF in mouse brain) that colocalize with poly(A) signal (red) in splicing speckles. **(3)** 3D rotation of an H4 cells expressing 0N4R tau-YFP showing intranuclear tau aggregates.

**Supplemental figure 1: FRAP of nuclear and cytosolic tau aggregates in HEK293 tau biosensor cells and additional examples of tau aggregates containing poly(A) RNA. (A)** H4 neuroglioma cells expressing 0N4R tau-YFP that contain nuclear tau aggregates. **(B, C)** FRAP of nuclear and cytosolic tau aggregates in HEK293 cells. 405 nm laser was used to photobleach and fluorescence intensity was measured in the 488cm channel (n=5). Arrow in example images shows bleached area. **(D)** FISH for Poly(A) RNA in HEK293 cell nuclear and cytosolic tau aggregates. Bar graphs show quantification of poly(A) RNA fluorescence intensity within nuclear and cytosolic tau aggregates in relation to bulk cytosol and nucleoplasm (n = 35 aggregates). White pixels in Coloc image show pixels above the Costes determined thresholds in 2D intensity plots (PCC = Pearson correlation coefficient and tM1= thresholded manders colocalization (% of tau pixels above threshold that colocalize with poly(A) pixels above threshold)). Error bars represent 95% confidence intervals and significance was determined using an unpaired two-tailed t-test. **(E)** Additional examples from B6 non transgenic and P301S mice (Tg2541) showing cytosolic tau aggregates (Ex. 1) and nuclear tau aggregates (Ex. 2, 3) contain RNA.

**Supplemental figure 2: Additional data on HEK293 and mouse brain tau aggregate isolation. (A)** Schematic showing isolation of HEK293 tau aggregates. HEK293 cells transfected with either WT or P301S mouse brain homogenate were lysed, and the cell lysate was stained with the RNA dye, SYTO17, showing the presence of RNA. The cell lysate was then run through a fluorescence activated particle sorter and gates were set to sort fluorescent particles in the P301S but not WT sample. Sorted fractions were examined by fluorescence microscopy to ensure particles were sorted. **(B)** Agilent Tapestation traces and QuBit readings of RNA isolated from sorted tau aggregates. **(C)** Flowcytometry scatter plots showing fluorescence and side scatter of particles for each replicate from the WT and P301S cell lysate (left column). The sorted and waste fractions were then re-run through the flow cytometer to ensure the particles were being sorted (right column). **(D)** Analysis of mRNAs two-fold enriched in tau aggregates from HEK293 cells. Genes from the following groups were highlighted: Voltage-gated calcium channel proteins, histone proteins, centrosomal proteins, and splicing related proteins. **(E)** Cellular component gene ontology of the mRNAs that are two-fold enriched in tau aggregates. **(F, G)** Analysis of multicopy gene families using RepEnrich showing enrichment of specific snoRNA repeats (U8, U17, and U3), snRNA repeats (U1 and U2), tRNA species, and Alu elements. The gene family color scheme is shared between F and G. **(H)** Schematic of tau aggregate isolation from mouse brain. Fractions that were taken for RNA isolation and sequencing are highlighted in Red. **(I)** Western blot of fractions from tau aggregate isolation showing P2 fraction (sarkosyl insoluble fraction) enriches for insoluble tau that is present in the P301L mice and not the WT mice. **(J)** Gene type enrichment in the P301L and WT sarkosyl insoluble samples. Fold change for each gene type was calculated by dividing the percentage of total FPKM made up by each gene type in the sarkosyl insoluble fraction (P2) by the percentage of total FPKM made up by each gene type in the total RNA. The numbers below the gene type names indicate the P301L/WT enrichment. Note the absence of snRNA enrichment in the sarkosyl insoluble fraction. **(K)** Mouse vs HEK293 gene type enrichment. **(L)** Scatter plot of mRNA enrichment in mouse brain tau aggregates in P301L and WT mice. Enrichment scores were calculated by dividing the insoluble tau IP FPKM by the total RNA FPKM for each replicate. mRNAs in red are two-fold enriched and mRNAs in blue are two-fold depleted from the P301L sarkosyl insoluble tau IP. Genes with fewer than 5 FPKM were removed from analysis due to low coverage. Two groups of mRNAs were highlighted: centrosomal proteins and splicing related mRNAs.

**Supplemental figure 3: Additional FISH for RNAs in HEK293 tau biosensor cells. (A, B)** FISH of the enriched RNU6ATAC and snoRA73B shows overlapping fluorescence intensity in nuclear tau aggregates. **(C)** FISH for enriched multicopy Alu RNAs. Quantification shows enrichment in to nuclear and cytosolic tau aggregates, with greater enrichment into cytosolic tau aggregates relative to nuclear aggregates. Bar graphs show quantification of FISH fluorescence intensity within nuclear and cytosolic tau aggregates in relation to bulk cytosol and nucleoplasm (n = 20 aggregates). Significance was determined using an unpaired two-tailed t-test. White pixels in Coloc images show pixels above the Costes determined thresholds in 2D intensity plots (PCC = Pearson correlation coefficient and tM1= thresholded manders colocalization (% of tau pixels above threshold that colocalize with Alu FISH pixels above threshold)). **(D)** FISH for the depleted mRNA, CENPQ, reveals lack of enrichment in nuclear and cytosolic tau aggregates.

**Supplemental figure 4: Additional HEK293 tau biosensor cell data supplementing figure 5. (A)** Images of HEK293 biosensor cells expressing SRRM2_FL-halo showing colocalization between nuclear and cytosolic tau aggregates. White pixels in Coloc images show pixels above the Costes determined thresholds in 2D intensity plots (PCC = Pearson correlation coefficient and tM1= thresholded manders colocalization (% of tau pixels above threshold that colocalize with SRRM2_FL-halo pixels above threshold)). **(B, C)** IF for two nucleolar proteins, Fibrillarin and RPF1, showing lack of colocalization between tau aggregates and the nucleolus. Costes method was unable to identify a positive correlation to determine thresholds. **(D)** Analysis of the % of cytosolic tau aggregates that colocalize with SRRM2 in HEK293 tau biosensor cells. 165 cells in 5 images were scored by hand as either colocalized or not colocalized. **(E)** Analysis of SRRM2 fluorescence intensity within cytosolic and nuclear tau aggregates relative to bulk cytosol and nucleoplasm (n = 20 measurements). Significance was determined using an unpaired two-tailed t-test. **(F)** Tau aggregates form in both the nucleus and the cytosol with and without lipofectamine 2000 as a transfection reagent showing that SRRM2 mislocalization to cytosolic tau aggregates is not dependent on lipofectamine. **(G, H)** Analysis of whether SRRM2 mislocalization disrupts the formation of SON positive splicing speckles. Cells outlined in white are cells that have SRRM2 mislocalized to the cytosol and by visual inspection and quantification of the average SON intensity, have no change in splicing speckles due to the mislocalization of SRRM2. Error bare represent 95% confidence intervals.

**Supplemental Figure 5: Additional data to support** **Fig. 5. (A)** Images of proteins examined for their association with tau aggregates in Fig 5 A, B. **(B)** Disorder prediction of SRRM2 by IUPred2 showing the relative ordered nature of the N-terminus and the disorder of the C-terminus. The cwf21domain and the location of the SRRM2_ dIDR-halo truncation are shown. **(C, D)** Schematic of the halo tagged SRRM2 fusion proteins made using CRISPaint and gel showing JF647 conjugated to halo fusion constructs running at the appropriate sizes

**Supplemental Figure 6: Additional splicing analysis data from cells with and without tau aggregates. (A)** Example of intron retention between exon 8 and 9 in ATF3 identified by MAJIQ at ΔPSI threshold of 0.1 and confidence threshold of 0.95. **(B)** Integrated genome viewer image showing raw read counts for each sequencing replicate. Intron retention can be seen between exon 8 and 9. **(C)** Gene ontology of the genes containing significantly retained introns in cells with tau aggregates (FDR < 0.05).

**Supplemental figure 7: Additional *in vivo* data from mouse and human tauopathies. (A)** IF and FISH showing p-tau (S422) positive tau aggregates colocalized with SRRM2 (SC-35) and poly(A) RNA in the nucleus of Tg2541 mouse hindbrain. White pixels in Coloc images show pixels above the Costes determined thresholds in 2D intensity plots (PCC = Pearson correlation coefficient and tM1= thresholded manders colocalization (% of tau pixels above threshold that colocalize with red pixels above threshold)). Y-axis rotation shows that the pTau(S422) foci being interrogated in the zoomed images are within the nucleus rather than above or below the nucleus. **(B)** IHC in human brain showing cytosolic inclusions of SRRM2 in AD and FTLD patient brains, but not in control brains. (Patient demographics for this figure and Figure 6B can be viewed in Supplemental Table 3).

## METHODS

### Tauopathy mouse models

Animals were maintained in a facility accredited by the Association for Assessment and Accreditation of Laboratory Animal Care in accord with the Guide for the Care and Use of Laboratory Animals. All procedures were approved by the University of California, San Francisco, Institutional Animal Care and Use Committee. Animals were maintained under standard environmental conditions, with a cycle of 12 hours light and 12 hours dark and free access to food and water.

For tau seeding experiments in HEK293 tau biosensor cells, the following mice were used: homozygous B6-Tg(Thy1-MAPT*P301S)2541 mice (referred to as Tg2541 or P301S tau mice in cellular seeding experiments) and FvBB6F1-Tg(Camk2a-tTa),(tetO-MAPT*wt)21221 (referred to as rTg21221 or WT tau mice in cellular seeding experiments). Mice were euthanized when the P301S tau mice developed spontaneous pathology (6-7 months). Collected brains were homogenized to 10% (wt/vol) in DPBS, aliquoted, and frozen at −80°C.

For tau isolation and sequencing of tau aggregates, the following mice were used: FvBB6F1-Tg(Camk2a-tTA)1Mmay, (tet)-tdTomato-Syp/EGFP)1.1Luo/J,(tetO-MAPT*P301L)4510 (referred to as rTg4510 or P301L mice in sequencing experiments) and FvBB6F1-Tg(Camk2a-tTa),(tetO-MAPT*wt)21221 (referred to as rTg21221 or WT tau mice in sequencing experiments).

For IF and FISH experiments, the following mice were used: homozygous B6-Tg(Thy1-MAPT*P301S)2541 mice (referred to as Tg2541 or P301S tau mice in IF and FISH experiments) and C57BL/B6 non transgenic mice (referred to as WT in IF and FISH experiments) were used as a control.

### Clarification of brain homogenate for tau aggregate seeding in HEK293 cells

10% brain homogenate from Tg2541 or WT mice was centrifuged at 500 x g for 5 minutes, the supernatant was transferred to a new tube and centrifuged again at 1,000 x g for 5 minutes. The supernatant was again transferred to a new tube and the protein concentration was measured using bicinchoninic acid assay (BCA), and diluted in DPBS to 1 mg/mL for transfection into HEK293 tau biosensor cells.

### PTA precipitation from brain homogenate for tau aggregate seeding in H4 biosensor cells

PTA precipitation of tau aggregates from mouse brain was performed as described (Woerman *et al,* 2016). 10% brain homogenate was incubated in final concentrations of 2% sarkosyl (Sigma, 61747) and 0.5% benzonase (Sigma, E1014-25KU) with constant agitation at 37°C for 2 hours. Sodium PTA (Sigma, P6395) was made in ultrapure MilliQ H_2_O and the pH was adjusted to 7.0. PTA was added to the samples to a final concentration of 2%, and samples were then incubated shaking at 37°C overnight. The samples were centrifuged at 13,200 × g at room temperature for 30 minutes, and the supernatant was removed. The resulting pellet was resuspended in 2% sarkosyl/PBS and 2% PTA. The sample was again incubated shaking at 37°C for 2 hours before a second centrifugation as above. The supernatant was again removed, and the pellet was resuspended in 1X PBS to 10% of the initial starting volume. This suspension was incubated using 1μL/well with Lipofectamine 2000 and OptiMEM at room temperature for at least 1.5 hours prior to infecting cells.

### Cell culture and tau seeding of H4 biosensor cells

H4 cells (ATCC, HTB-148) stably expressing the *pIRESpuro3* vector (Clontech) containing a codon-optimized 0N4R MAPT gene with the P301S point mutation and tagged with YFP were cultured in Dulbecco’s Modified Eagle’s Medium (DMEM) supplemented with 10% fetal bovine serum (FBS) and 0.2% penicillin-streptomycin, and maintained in incubators set to 37°C with 5% carbon dioxide. Cells were plated in a 12-well glass-bottomed dish at 1x10^5^ cells/well and allowed to settle for a minimum of 2 hours prior to infection with PTA-precipitated tau prions from Tg2541 mouse brain.

### Cell culture and tau seeding in HEK293 biosensor cells

HEK293 biosensor cells stably expressing the 4R RD of tau with the P301S mutation were purchased from ATCC (CRL-3275) (previously described in (Holmes et al., 2014)). Cells were seeded at 2.5 x 10^5^ cells/mL in 500uL of DMEM with 10% FBS and 0.2% penicillin-streptomycin antibiotics on PDL coated glass coverslips in a 24-well tissue culture treated plate (Corning 3526) and allowed to grow overnight in incubators set to 37°C with 5% carbon dioxide. The next day, 7ug of 1 mg/mL clarified P301L tau or WT tau mouse brain homogenate was mixed with 6uL of Lipofectamine 2000 and brought up to 100uL in PBS and allowed to sit at room temperature for 1.5 hours. The mixture was then added to 300uL of DMEM without FBS or antibiotics and mixed by pipetting. 50uL of this mixture was added to each well of a 24 well plate and allowed to incubate at 37 °C for 24 hours. Tau aggregate formation was monitored using a fluorescence microscope with a 488nm filter.

### Generation of Lentiviral particles

As previously described (Burke et al., 2019), HEK293T cells (T25 Flask at 80% confluence) were co-transfected with 1ug of pLenti-SRSF2-mCherry-blasticydin, 1ug of pVSV-G, 1ug of pRSV-Rev, and 1ug of pMDLg-pRRe using 16uL of lipofectamine 2000. Medium was replaced 6 hours post-transfection. Medium was then collected at 24- and 48-hours post-transfection and filter sterilized with a 0.45-um filter.

### Generating SRSF2-mCherry cells via lenti-virus

HEK293 biosensor cells were seeded in a T-25 flask. When 80% confluent, the cells were incubated for 1 hour with 1mL of SRSF2-mCherry-blasticydin lentiviral particles containing 10-ug of polybrene with periodic rocking. 4mL of normal medium was then added to the flask and incubated for 24 hours. Normal medium was then aspirated and replaced with selective medium containing 10-ug/mL of Blasticidine S hydrochloride (Sigma-Aldrich). Selective medium was changed every three days. After one-week, selective medium was replaced with normal growth medium. Expression of SRSF2-mCherry was confirmed by fluorescence microscopy.

### Generating halo tagged SRRM2 cells using CRISPaint

HEK293 biosensor cells were seeded in a 6 well plate. As previously described (Ilik et al., 2020; Schmid-Burgk et al., 2016) when 80% confluent, cells were transfected with 1 ug of pCRIPaint-HaloTag-PuroR plasmid, 0.5 ug of PX458-CAS9 targeting plasmid, 0.5 ug of of pCAS9-mCherry-Frame_selector plasmid. After 24 hours, cells were selected using 2 ug/mL puromycin for 48 hours to enrich for edited cells. To label the halo constructs, JF646 was added to growth media at 10 nM overnight prior to cell lysis for gel analysis or fixation for imaging.

### Immunofluorescence (IF) in HEK293 and H4 cells

Cells were fixed in 4% FPA for 10 minutes, washed 3X with DPBS, permeabilized in 0.1% Triton X-100 (Fisher BP151-100) for 5 minutes, washed 3x with PBS, and blocked with 5% bovine serum albumin (BSA) for 1 hour. Primary antibodies were diluted to desired concentration in 5% BSA and incubated overnight at 4 deg. Slides were washed 3x with DPBS and secondary antibodies were added at appropriate dilution in 5% BSA and allowed to incubate at room temperature for 1 hour. Slides were washed 2x with DPBS and then incubated in DAPI diluted in PBS (1ug/mL) for 5 minutes at RT, washed 1X with DPBS and then mounted on microscope slides with Prolong glass antifade mountant.

### Fluorescence in-situ hybridization (FISH)

As previously described (Khong et al., 2017), cells were fixed in 4% PFA for 10 minutes, washed 3x with PBS, permeabilized in 70% ethanol for 1 hour at 4 deg. Cells were then incubated in a wash buffer consisting of 2X nuclease free SSC and 10% deionized formamide (Calbiochem 4610) for 5 minutes at room temperature. The FISH probes were diluted to desired concentration in 100uL of hybridization buffer (2X nuclease-free SSC, 10% deionized formamide, 10% dextran sulfate) and spotted onto parafilm in a hybridization chamber (10cm cell culture dish lined with wet paper towels and covered with parafilm). Coverslips were then inverted onto the droplet of hybridization buffer contain the FISH probes and incubated at 37°C overnight. The slides were then transferred back to a 24 well plate and 500uL of 2X nuclease free SSC with 10% deionized formamide was added for 30 minutes at 37°C. Cells were then incubated in DPBS with 1ug/mL DAPI at room temperature for 5 minutes, washed with 2X nuclease free SSC and incubated at room temperature for 5 minutes. Coverslips were mounted on microscope slides with ProLong Glass Antifade Mountant (ThermoFisher, P36980) and allowed to cure overnight at room temperature.

minutes.

### Image analysis

To quantify FISH intensity within nuclear and cytosolic tau aggregates, ImageJ’s freehand selection tool was used to draw perimeters around aggregates and in regions of bulk cytosol and nucleus. The average FISH intensity was measured within the selections and used to compare enrichment of RNAs.

To quantify the percentage of SRRM2 in the cytosol of cells, CellProfiler was used. Nuclei were identified using object detection with a typical diameter between 50-200 pixels for HEK293 and H4 cells, which were then used as a mask to quantify nuclear and cytosolic SRRM2 intensity. The percentage of total SRRM2 intensity in the cytosol was calculated by dividing the cytosolic SRRM2 intensity by the sum of the nuclear and cytosolic intensities per image.

To quantify enrichment scores of various proteins of interest (POIs) in tau aggregates, 25 images were taken in a 5x5 panel of each slide using a 40x air objective on a Nikon Spinning Disc Confocal microscope. Ilastik was used to create the following segmentation masks: cytosolic tau aggregates, nuclear tau aggregates, nucleus, cytosol, and background. The RGB images and segmentation masks were fed into CellProfiler, which was used to quantify the average POI intensity within tau aggregates and the average POI intensity within the corresponding compartment. Enrichment was defined as the ratio of POI intensity within the tau aggregate to the POI intensity within the corresponding compartment.

### Fluorescent labeling of oligonucleotides for FISH

As previously described (Gaspar et al., 2017), DNA oligonucleotides were labeled with ddUTP-Cy5 fluorophores using terminal deoxynucleotidyl transferase (TdT). DNA oligonucleotides were designed to be antisense to the target of interested with the following specifications: 18-22 nucleotides in length and a minimum of 2 nucleotide spacing between probes. 20uM of DNA oligonucleotides were mixed with 120uM of 5-Propargylamino-ddUTP-Cy5, 10 units of TdT, and 1X TdT buffer and incubated at 37°C for 16 hours. Following incubation, oligonucleotides were precipitated in 80% ethanol with 60mM Na-acetate at −80°C for 20 minutes. The oligonucleotides were pelleted by centrifugation at 16,000g for 20 minutes at 4°C, washed with 80% ethanol 2x, air dried, and brought up in 20uL of nuclease free H20. If necessary, a further round of purification can be performed with the Zymo Oligo Clean and Concentrate spin-column kit (Zymo D4060). Labeled probe concentration was measured via NanoDrop OneC UV-Vis Spectrophotometer (Thermo Scientific 840-274200).

### Tau aggregate isolation from HEK293 cells via centrifugation and flow cytometry

HEK293 biosensor cells were grown to 70-80% confluency in 245 mm square tissue culture treated dishes (Corning 07-200-599) in 50mL of DMEM (one plate per biologic replicate). 200 ug of WT or P301S tau clarified mouse brain homogenate was transfected per dish using lipofectamine 2000 and incubated at 37°C for 24 hours. Tau aggregation was monitored using the Evos M500 Imaging System with a GFP filter. Cells were harvested by scraping, centrifuged at 200 rcf, snap frozen in liquid nitrogen, and stored at −80°C.

The cell pellet was thawed on ice for 5 minutes and resuspended in 6mL of high salt, high sucrose buffer containing RNase Inhibitors (10mM Tris-HCl pH 7.4, 0.8M NaCl, 1 mM EGTA, 10% sucrose, 0.5% NP40, Complete ultra-protease inhibitor, PhosStop Phosphatase inhibitor, 1:1500 RNasein, 1:300 Ribolock, 1:60 turbo DNAse). Cell lysate was passed through a 25 G needle 3x to homogenize and 100uL of sample was taken to extract total RNA.

Large tau complexes were pelleted by centrifugation at 21,000g for 20 minutes at 4 °C, the pellet was brought up in high salt/high sucrose buffer, passed through a 27G needle, and centrifuged at 10,000g for 10 minutes at 4 °C. The pellet was brought up in 1mL of DPBS and centrifuged at 500g for 5 minutes at 4 °C to pellet large cellular debris. The supernatant (S3, enriched tau fraction) was taken and spotted onto a microscope slide for fluorescent imaging of tau aggregates in solution.

A BD Biosciences FACSAria Fusion flow cytometer was used to sort tau aggregates by fluorescence and size. The sheath fluid was changed to PBS, flow rate was set to 1.2, and threshold rate was set to <200 events/second. Gates were set on side scatter-H and 488 fluorescence such that WT transfected S3 fractions had <1% of particles in sorted fraction and P301S transfected S3 fractions had >30% in the sorted fraction. Roughly 1 million particles were sorted for each sample. To ensure the flow cytometer was sorting particles properly, the sorted fraction was visually inspected by fluorescence microscopy and the sorted and waste fractions were run back through the flow cytometer.

To denature tau aggregates and extract RNA, the sorted fractions were brought up in Proteinase K buffer (2M Urea, 100ug/mL proteinase K, and 3mM DTT) and incubated at room temperature for 15 minutes. Guanadinium HCl was added to a final concentration of 5M and incubated at room temperature for 30 minutes. RNA was then extracted with TRIzol LS reagent (ThermoFisher 10296010). RNA concentrations were measured by QuBit RNA HS Assay kit (ThermoFisher Q328521) and Agilent 4200 TapeStation using the High Sensitivity RNA ScreenTape (Agilent 5067-5579). All samples except for the WT transfected sorted fraction yielded sufficient RNA to prepare sequencing libraries (Fig S2B for Tapestation data). RNA sequencing libraries were then prepared from total RNA and tau aggregate associated RNA from HEK293 biosensor cells using the Roche KAPA RNA HyperPrep Kit with RiboErase (Kapa KK8560) and sequenced on an Illumina NextSeq sequencer at the University of Colorado, Boulder BioFrontiers Sequencing Core.

### Isolation of tau aggregates from mouse brain

Brains were harvested from two Tg21221 (WT 0N4R human tau mouse brains) and two rTg4510 (P301L 0N4R human tau) mice and snap frozen in liquid nitrogen. Samples were thawed on ice and weighed. The brain tissue was then homogenized on ice using a dounce homogenizer and diluted to 5 mL/g in homogenization buffer with RNase inhibitors (10mM Tris-HCl pH7.4, 0.8M NaCl, 1mM EGTA, 10% sucrose, 1X Roche protease inhibitor, 1:40 promega RNasein). Aliquots were stored at −80 C.

To extract total RNA, 50uL of brain homogenate was incubated for 2 hours at room temperature in proteinase K buffer (2% SDS, 4M Urea, 10mM Tris-HCl pH4.54, 100ug/mL Proteinase K). 400uL of Urea buffer (60mM Tris-HCl pH 8.5, 8M Urea, 2% SDS) was then added and incubated for 30 minutes at room temperature. RNA was extracted from one half of this reaction using TRIzol LS solution and the other half was frozen at −80°C.

900uL of frozen brain homogenate was thawed on ice and 100uL of 10% (w/v) sarkosyl solution was added and incubated on ice for 15 minutes. Homogenate was then passed through a 25G and 27G syringe. Protein concentrations were measured by QuBit Protein Assay Kit (Thermo Fisher, Q33211) and sarkosyl buffer (50mM HEPES pH 7.2, 250 mM sucrose, 1mM EDTA, 1% w/v sarkosyl, 0.5 M NaCl) was added to reach a final concentration of 10 mg/mL. 500uL of each sample was transferred into an untracentrifuge tube (Beckman Coulter, 349623) and centrifuged at 180,000g for 30 min at 4 deg in a Beckman Coulter Optima MAX-XP Ultracentrifuge. The supernatant (S1 fraction) was removed and stored at −80°C. The pellet was then brought up in 500uL of sarkosyl buffer and run through a 25G needle to homogenize. The sample was then centrifuged at 180,000g for 30 min at 4°C and the supernatant (S2) was removed and stored at −80°C.

For the sarkosyl insoluble RNA sequencing, P2 pellet was brought up in 100uL of proteinase K buffer and incubated at RT for 2 hours at RT. To further solubilize the sample, 400uL of urea buffer was added and incubated at RT for 30 minutes. The sample was then split in two and RNA was extracted from one half (250uL) using Trizol LS solution the other half was frozen at −80°C.

For the Tau IP, the P2 fraction was brought up in 400uL of PBS and protein concentrations were measured using the QuBit Protein Assay Kit (Thermo Fisher, Q33211). Samples were precleared with 15mg of DEPC treated (to inactivate RNAse) protein A dynabeads at room temperature for 45 minutes at RT on rotator. While preclearing, Tau12 and IgG antibodies were conjugated to 50uL (1.5mg) of protein A dynabeads for 40 minutes on rotator. Following preclear step, the sample was split into two fractions (one for the Tau12 IP and one for the IgG IP). Dynabeads with conjugated antibody were washed with PBS, brought up in 50uL of PBS and added to sample. IP was carried out on rotator at room temperature for 40 minutes. Sample was then washed 3x with PBS and 100uL of proteinase K buffer was added to the beads and incubated at RT for 2 hours. 400uL of urea buffer was added to beads and incubated for 30 minutes to further denature. Samples were then split into two 250uL fractions, one was Trizol extracted. RNA concentrations were then measured by QuBit RNA HS Assay kit (ThermoFisher Q328521) and Agilent 4200 TapeStation using the High Sensitivity RNA ScreenTape (Agilent 5067-5579). IgG IP did not pull down any RNA. RNA sequencing libraries were prepared using the Nugen Ovation SoLo RNA-Seq System, Mouse (Nugen 0501-32) and sequenced on an Illumina NovaSeq sequencer at the University of Colorado, Anschutz Genomics and Microarray Core.

### Analysis of RNA sequencing data

Following sequencing, quality of sequencing reads were assessed using FASTQC version 0.11.5, Illumina TruSeq3 adapters and low quality reads were trimmed off using Trimmomatic version 0.36 (Bolger et al., 2014). Reads that aligned uniquely to the ribosome, yeast, or bacteria were then filtered out using FastQ Screen (Wingett and Andrews, 2018). Reads were aligned using the Spliced Transcripts Alignment to a Reference (STAR) aligner version 2.6.0 (Dobin et al., 2013) to either the Genome Reference Consortium Human Build 38 (GRCh38, acquired from NCBI) or the Genome Reference Consortium Mouse Build 38 (GRCm38, acquired from NCBI) depending on the species being analyzed. Adjusted p-values were calculated from raw read counts using DEseq2. Gene counts were used to calculate Fragments per kilobase per million read (FPKM) using transcript lengths retrieved from the Ensembl Biomart (Kinsella et al., 2011) and the following formula FPKM = (# of mapped fragments*10^3^*10^6^)/(transcript length in bp * total number of mapped fragments). For mouse sequencing, FPKM values were used to calculate enrichment scores for each biological replicate by dividing the Tau IP FPKM by the Total RNA FPKM for each replicate. Enrichment scores were then used to calculate average enrichment score for each gene and fold changes between P301L and WT mice. Gene type enrichment was determined by calculating the percentage of FPKM made up by each gene type. Repetitve elements were analyzed using a reference files acquired from repeatmasker (hg38 - Dec 2013 - RepeatMasker open-4.05 – Repeat Library 20140131) and RepEnrich (Criscione et al., 2014).

Splicing analysis: Following mapping of reads to GRCh38 using STAR, MAJIQ v2.1 was used with standard settings to quantify splicing changes. Voila was used to view results, generate splicing diagrams, and determine the relative percentage of each splicing type (only LSVs containing more than 10 reads were reported). To quantify reads mapping to introns, iREAD v0.8.5 was used along with an intron annotation file generated from ensemble v77. The read count output from iREAD was used as an input to DEseq2 for calling differential intron retention.

### Fluorescence recovery after photobleaching

HEK293 biosensor cells were seeded in DMEM supplemented with 10% FBS and 0.2% penicillin-streptomycin at 0.25x10^5^ cells/mL in Grenier Bio-One CELLview dishes with Glass Bottoms (Thomas Scientific, 07-000-235) and grown overnight at 37°C. The next day, clarified P301S tau brain homogenate was transfected and grown for 24 hours. A Nikon A1R Laser Scanning Confocal with environmental chamber was used to image the cells. A circular region within a tau aggregate was defined and bleached using a 405nm laser set to 100% laser power. For determining the recovery of tau within tau aggregates (Fig S1B-C), fluorescence intensity was measured continuously for 6 minutes and 30 seconds post bleaching (N=5). For determining the recovery of SRRM2_FL-halo within splicing speckles with and without tau aggregates, fluorescence intensity was measured continuously for 30 seconds (n=5). For determining the recovery of SRSF2-mCherry within splicing speckles with and without tau aggregates, fluorescence intensity was measured every 1 second for 30 seconds (n=5).

### Mouse brain RNA fluorescent *in situ* hybridization followed by immunofluorescent staining (RNA FISH-IF)

Control (B6/J) and Tg2541 animals at approximately 6 months of age were anesthetized for whole brain collection. The mouse brains were embedded in OCT compound (Sakura, 4583) and flash-frozen in chilled isopentane. Samples were sectioned at 12μm using a cryostat and mounted on glass slides. Samples were air-dried at room temperature for 20 minutes to ensure tissue adherence to slides, then fixed in cold 4% PFA/1X PBS for 15 minutes. Samples were washed 3 times in 1X PBS for 5 minutes/wash, followed by a wash in 1X SSC for 5 minutes. Samples were transferred into 0.1X citrate buffer (Sigma, C9999) for a gentle antigen retrieval at 60°C for 1 hour 15 minutes. The slides were allowed to cool for 15 minutes, then rinsed 3 times in 1X SSC for 5 minutes/wash. Samples were then dehydrated in a graded series of ethanol washes (50%, 70%, 90% and 100%) for 3 minutes/wash and air-dried for 10 minutes. A hydrophobic barrier was drawn around the tissue and samples were blocked in a pre-hybridization buffer of 3% normal goat serum (NGS)/4X SSC at 37°C for 1 hour in a humidified chamber. Oligo(dT) probe labeled with Quasar 570 or Quasar 670 (Stellaris) was added to hybridization buffer (Stellaris, SMF-HB1-10) and incubated at 65°C for 10 minutes followed by a cooling on ice for 2 minutes. Pre-hybridization buffer was removed, and samples were then incubated in probe/hybridization buffer at 37°C overnight.

The next day, samples were washed in a dilution series of pre-hybridization buffer (twice in 4X, then once in 2X, 1X, 0.1X) at 37°C for 10 minutes/wash. Samples were then blocked in 20% NGS/1X PBST (0.1% Tween-20) at room temperature for 1 hour and incubated at room temperature overnight in primary antibodies diluted at 1:250 in 10% NGS/1X PBST.

The next day, samples were washed 3 times in 1X PBST for 10 minutes/wash, then incubated in Alexa Fluor secondary antibodies diluted at 1:500 in 10% NGS/1X PBST for 2 hours at room temperature. Samples were washed 3 times in 1X PBST for 10 minutes/wash. To quench autofluorescence, samples were incubated in 0.1% Sudan Black B in 70% ethanol for 10 minutes, then rinsed briefly in fresh 70% ethanol and transferred into 1X PBS for 5 minutes. Coverslips were mounted onto the slides using Vectashield Vibrance antifade mounting medium with DAPI (Vector Laboratories H-1800) and slides were left to dry overnight prior to being imaged with a Leica SP8 confocal microscope.

### Human brain immunofluorescent staining

Human brain samples were provided by the Neurodegenerative Disease Brain Bank at the University of California, San Francisco, which receives funding support from NIH grants P01AG019724 and P50AG023501, the Consortium for Frontotemporal Dementia Research, and the Tau Consortium.

Formalin-fixed paraffin-embedded (FFPE) human brain samples from the left angular gyrus region of control individuals and patients diagnosed with corticobasal degeneration (CBD) were sliced at 8μm and mounted on glass slides. Samples were deparaffinized in a 60°C oven overnight, followed by two 10-minute xylene washes. Samples were then rehydrated in a graded series of ethanol washes (twice in 100% then once each in 90%, 70%, 50%) for 3 minutes/wash. Slides were then rinsed in cold ultrapure MilliQ H_2_O and transferred into 0.1X citrate buffer (Sigma, C9999) for antigen retrieval in an autoclave at 120°C for 5 minutes. Slides were allowed to cool for 15 minutes, then rinsed in 1X PBST (0.25% Triton X-100) for 15 minutes. A hydrophobic barrier was drawn around the tissue and samples were blocked in 20% normal goat serum (NGS)/1X PBST at room temperature for 1 hour in a humidified chamber. Samples were incubated at room temperature overnight in primary antibodies diluted at 1:250 in 10% NGS/1X PBST.

The next day, samples were washed 3 times in 1X PBST for 10 minutes/wash, then incubated in Alexa Fluor secondary antibodies diluted at 1:500 in 10% NGS/1X PBST for 2 hours at room temperature. Samples were washed 3 times in 1X PBST for 10 minutes/wash. To quench autofluorescence, samples were incubated in 0.1% Sudan Black B in 70% ethanol for 10 minutes, then rinsed briefly in fresh 70% ethanol and transferred into 1X PBS for 5 minutes. Samples were incubated for 10 minutes in 5μg/mL DAPI diluted in 1X PBS, then washed for 10 minutes in 1X PBS. Coverslips were mounted onto the slides using Vectashield Vibrance antifade mounting medium (Vector Laboratories H-1700) and slides were left to dry overnight prior to being imaged with a Leica SP8 confocal microscope.

### Human tissue samples for immunohistochemistry

AD, FTD and non-neurologic disease control post-mortem tissue samples were obtained from the University of Pittsburgh ALS Tissue Bank, the Barrow Neurological Institute ALS Tissue Bank, and the Target ALS Human Postmortem Tissue Core. All tissues samples were collected after informed consent from the subjects or by the subjects’ next of kin, complying with all relevant ethical regulations. The protocol and consent process were approved by the University of Pittsburgh Institutional Review Board (IRB) and the Dignity Health Institutional Review Board. Clinical diagnoses were made by board certified neuropathologists. Subject demographics are listed in Supplemental Table 3.

### Immunohistochemistry

Paraffin-embedded post-mortem frontal cortex tissue sections were used for this study. All sections were deparaffinized, rehydrated and antigen retrieval performed using Target Antigen Retrieval Solution, pH 9.0 (DAKO) for 20 min in a steamer. After cooling to room temperature, non-specific binding sites were blocked using Super Block (Scytek), supplemented with Avidin (Vector Labs). Primary antibodies used for immunohistochemistry were incubated overnight in Super Block with Biotin. Slides were then washed and incubated for 1 h in the appropriate biotinylated IgG secondary antibodies (1:200; Vector Labs) in Super Block. Slides were washed in PBS and immunostaining visualized using the Vectastain Elite ABC reagent (Vector Labs) and Vector Immpact NovaRED peroxidase substrate kit (Vector Labs). Slides were counterstained with hematoxylin (Sigma Aldrich) and pictures were captured using an OLYMPUS BX40 microscope equipped with a SebaCam camera.

### Antibody information

| **Antibody name** | **Manufacturer** | **Catalog #** |
| --- | --- | --- |
| Mouse anti-phospho tau S202, Thr205 | Invitrogen | MN1020 |
| Rabbit anti-pTau Thr205 | Invitrogen | 44-738G |
| Mouse anti-pTau (Ser202, Thr205) AT8 | Invitrogen | MN1020 |
| Mouse anti-SRRM2 (SC-35) | Abcam | Ab11826 |
| Rabbit anti-Pinin | Thermo Scientific | 18266-1-AP |
| Rabbit anti-SFPQ | Abcam | Ab177149 |
| Rabbit anti-ZC3H14 (MSUT2) | Sigma-Aldrich | HPA049798 |
| Rabbit anti-DDX39B | OriGene | TA890032 |
| Rabbit anti-DYRK1A | Abcam | ab65220 |
| Rabbit anti-SRSF2 | Abcam | Ab204916 |
| Rabbit anti-SNRNP70 (U1-70K) | Abcam | ab83306 |
| Rabbit anti-PABP | Abcam | ab21060 |
| Rabbit anti-TIA1 | Abcam | ab40693 |
| Rabbit anti-SRRM3 | Sigma-Aldrich | HPA019337 |
| Rabbit anti-SRSF1 (SF2) | Abcam | ab129108 |
| Mouse anti-G3BP | Abcam | ab56574 |
| Rabbit anti-SNRPA1 | Millipore Sigma | HPA045622 |
| Rabbit anti-PQBP1 | Millipore Sigma | HPA001880 |
| Rabbit anti-PNISR | Millipore Sigma | HPA038796 |
| Rabbit anti-RPF1 | Sigma-Aldrich | HPA024642 |
| Rabbit anti-U2AF1 | Sigma-Aldrich | HPA044833 |
| Rabbit anti-SON | Thermo Fisher | PA565108 |
| Rabbit anti-Fibrillarin | Invitrogen | MA1-22000 |
| Goat anti-mouse 647 | Abcam | Ab150115 |
| Goat anti-rabbit 647 | Abcam | Ab150079 |

### Fluorescence in-situ hybridization probe information

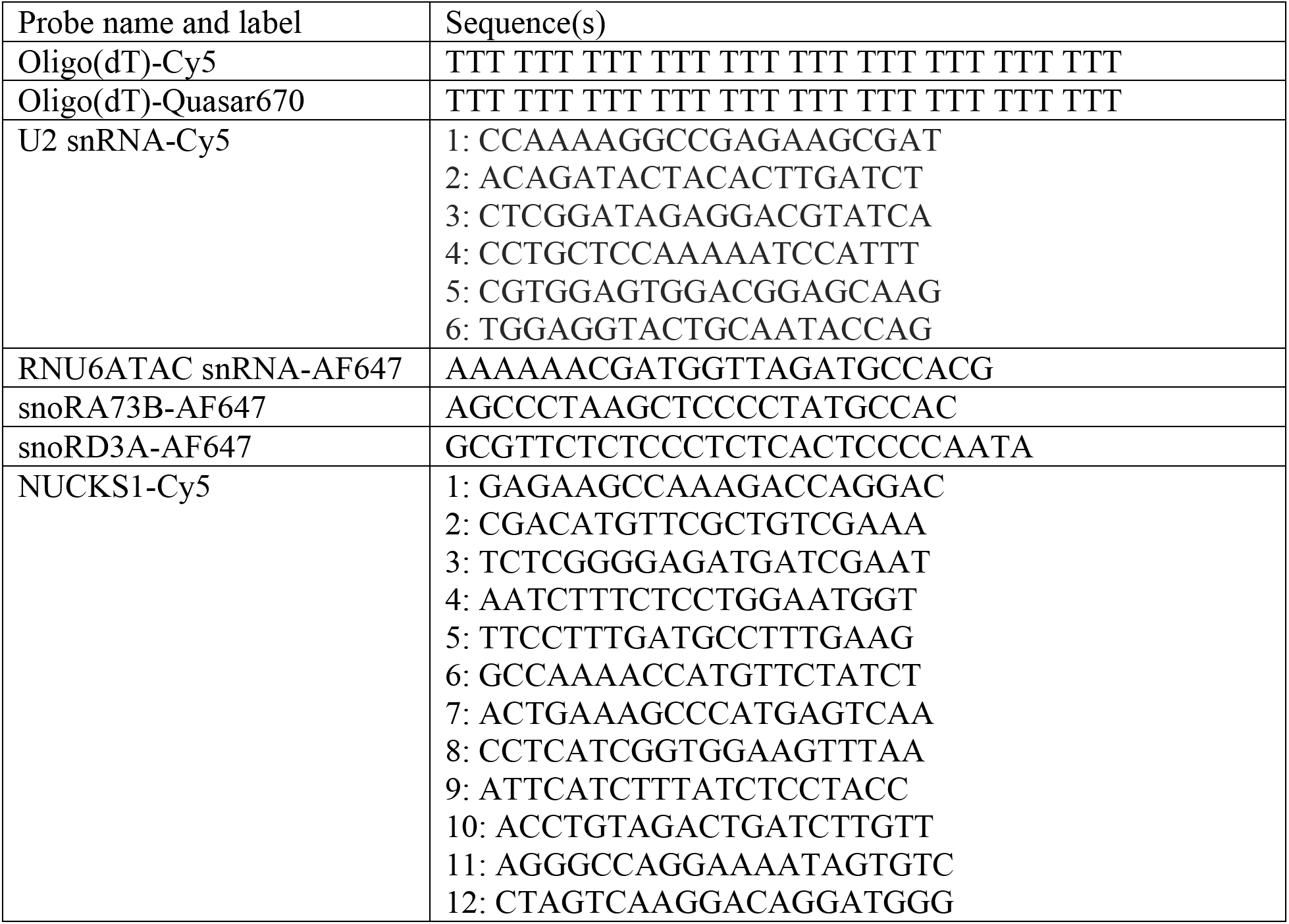
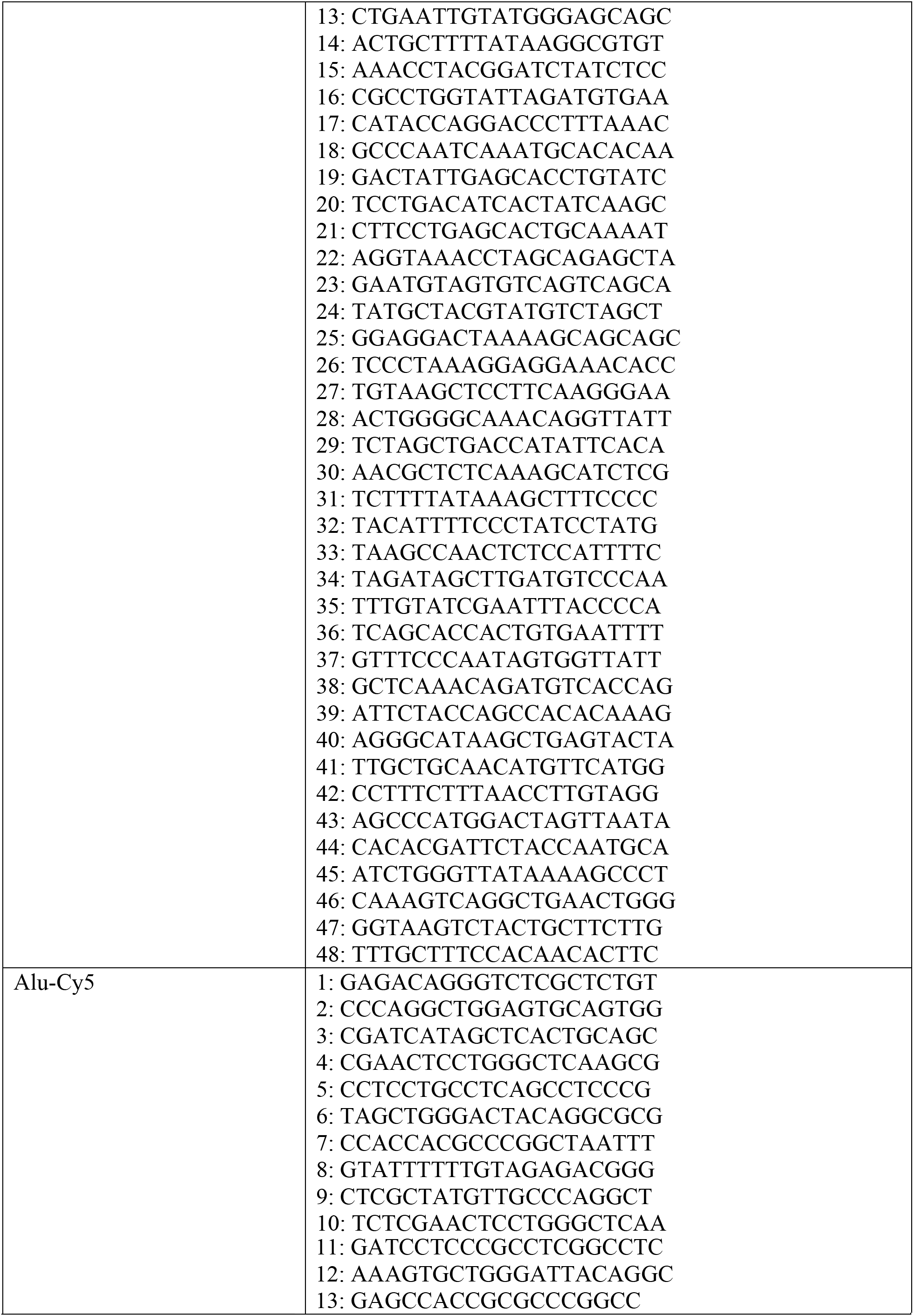

