## Supplemental Figures for "Tau aggregates are RNA-protein assemblies that mis-localize multiple nuclear speckle components"

**Supplemental Figure 1: FRAP of nuclear and cytoplasmic tau aggregates in HEK293 tau biosensor cells and additional examples of tau aggregates containing poly(A) RNA**

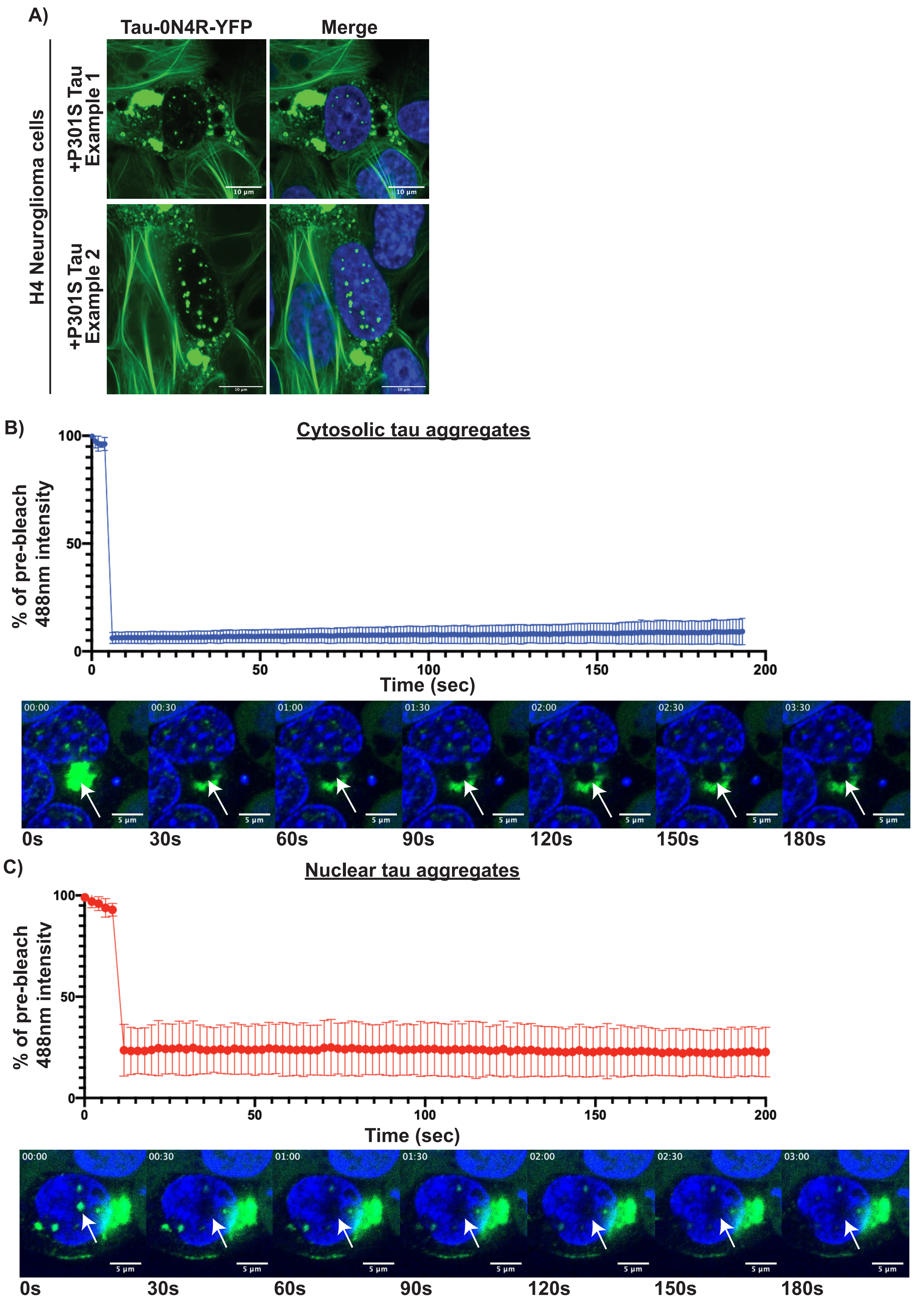

**Supplemental Figure 1 continued: Tau aggregates in HEK293 biosensor cells and mice contain poly(A) RNA**

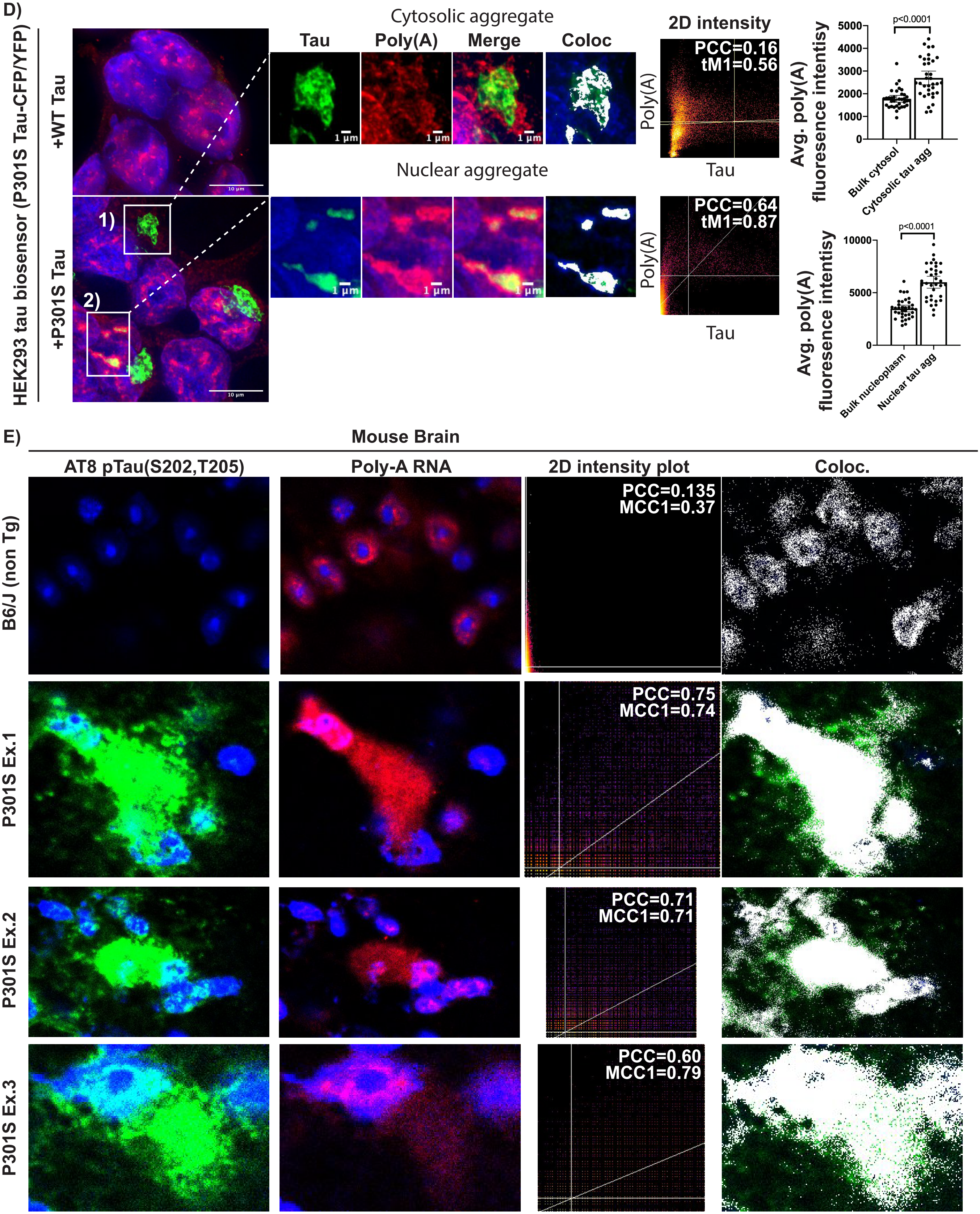

### Supplemental Figure 2: Additional data on HEK293 and mouse tau aggregate isolation and sequencing

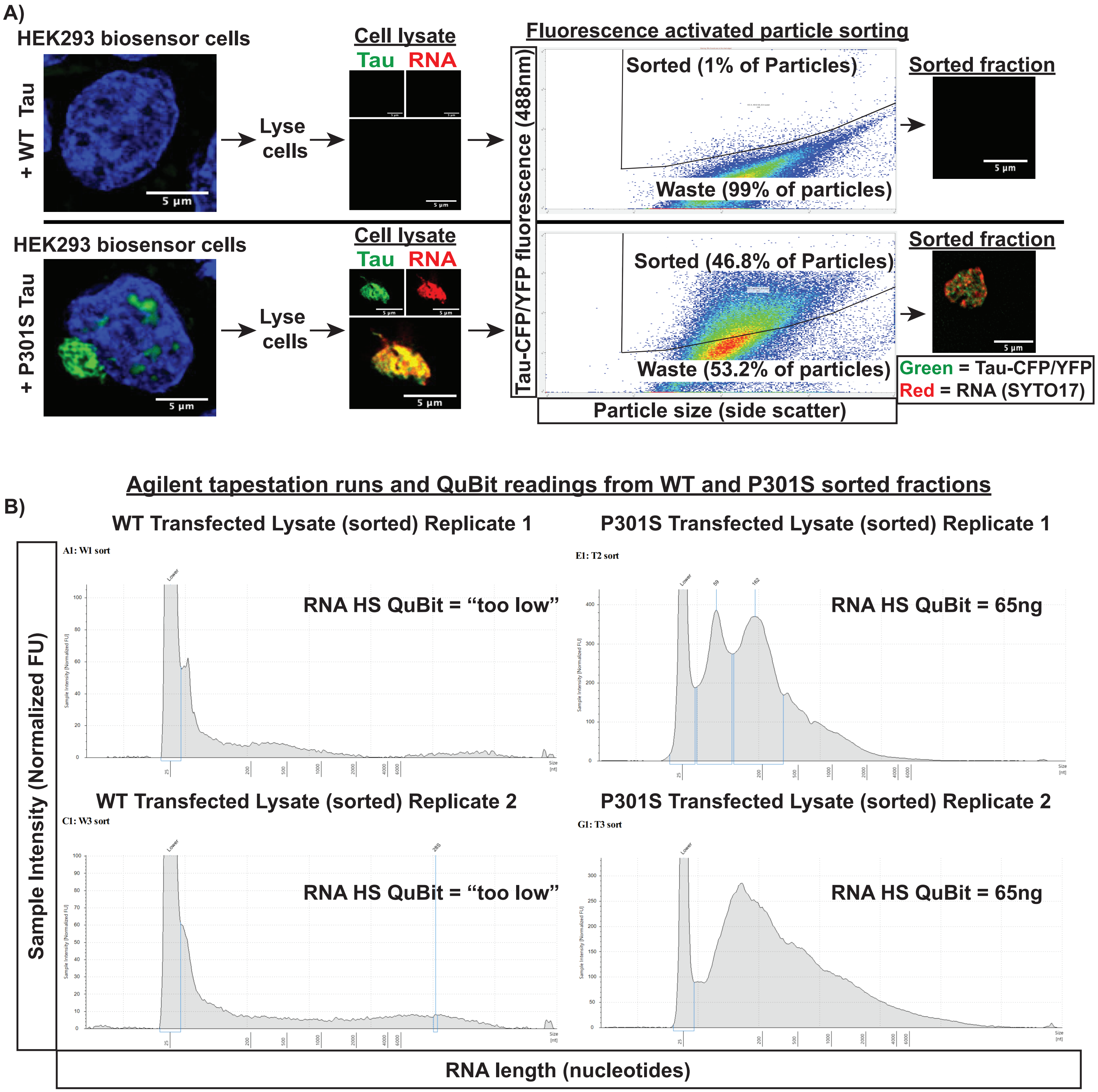

**Supplemental Figure 2 continued: Additional data on HEK293 and mouse tau aggregate isolation and sequencing**

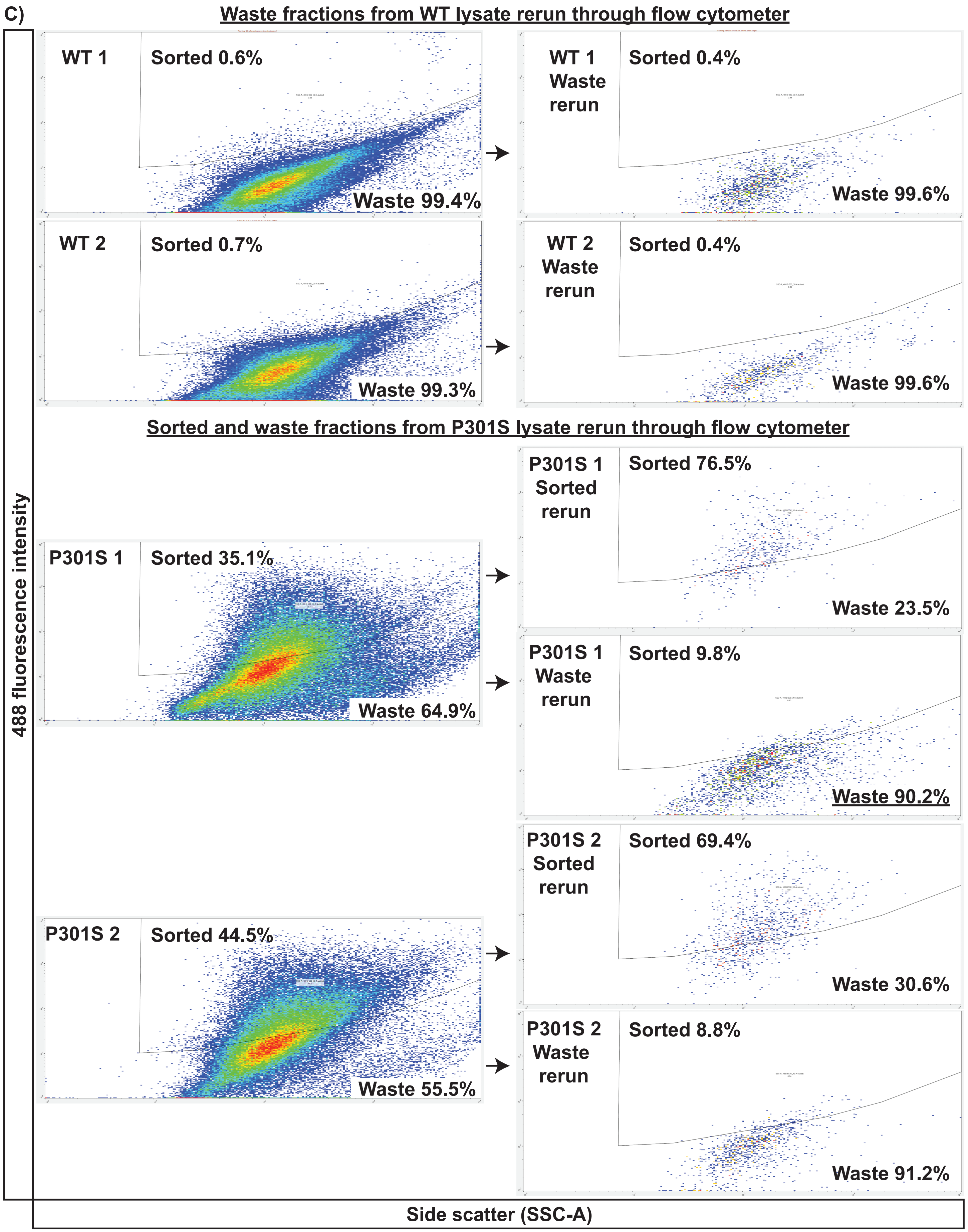

**Supplemental Figure 2 continued: Additional data on HEK293 and mouse tau aggregate isolation and sequencing**

D) **mRNAs significantly enriched in HEK293 tau aggregates**

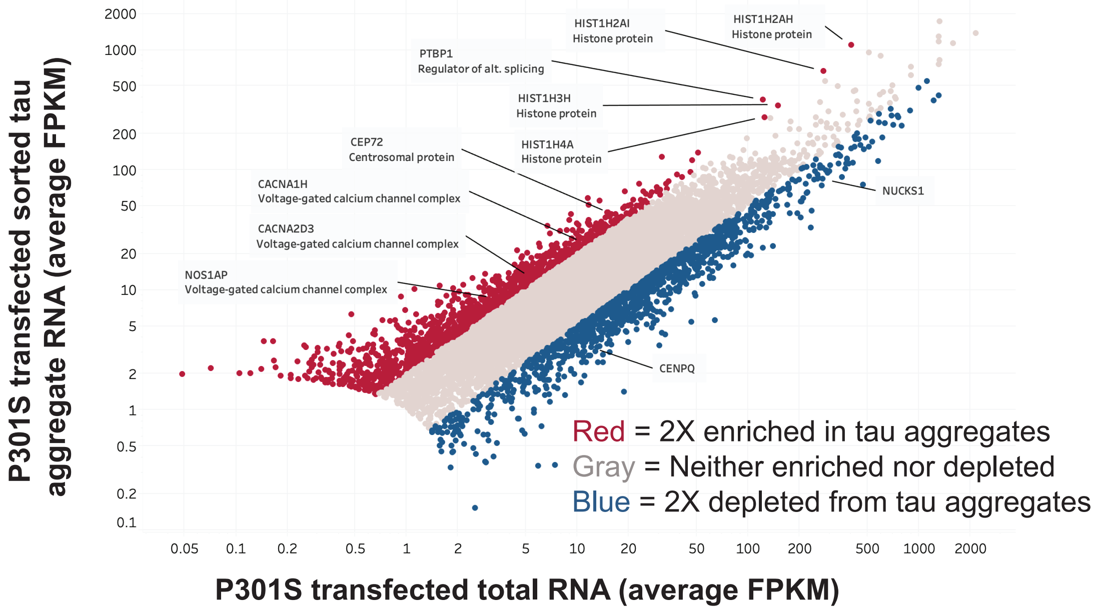


E) **Gene ontology of 2x enriched mRNAs - cellular component**

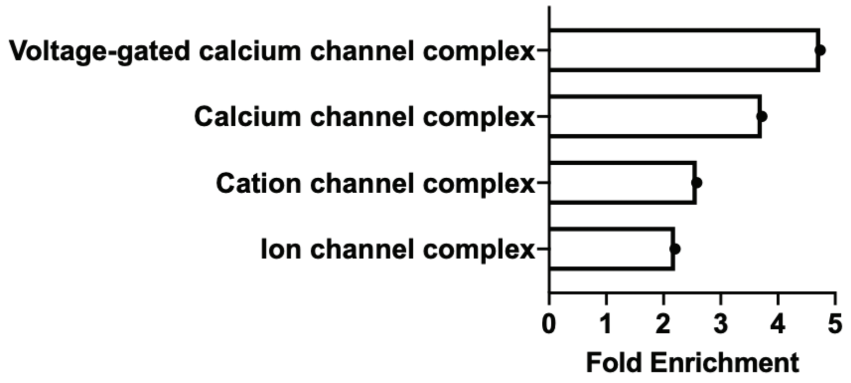


**Analysis of multicopy gene families in HEK293 tau biosensor tau aggregates using RepEnrich**

F)

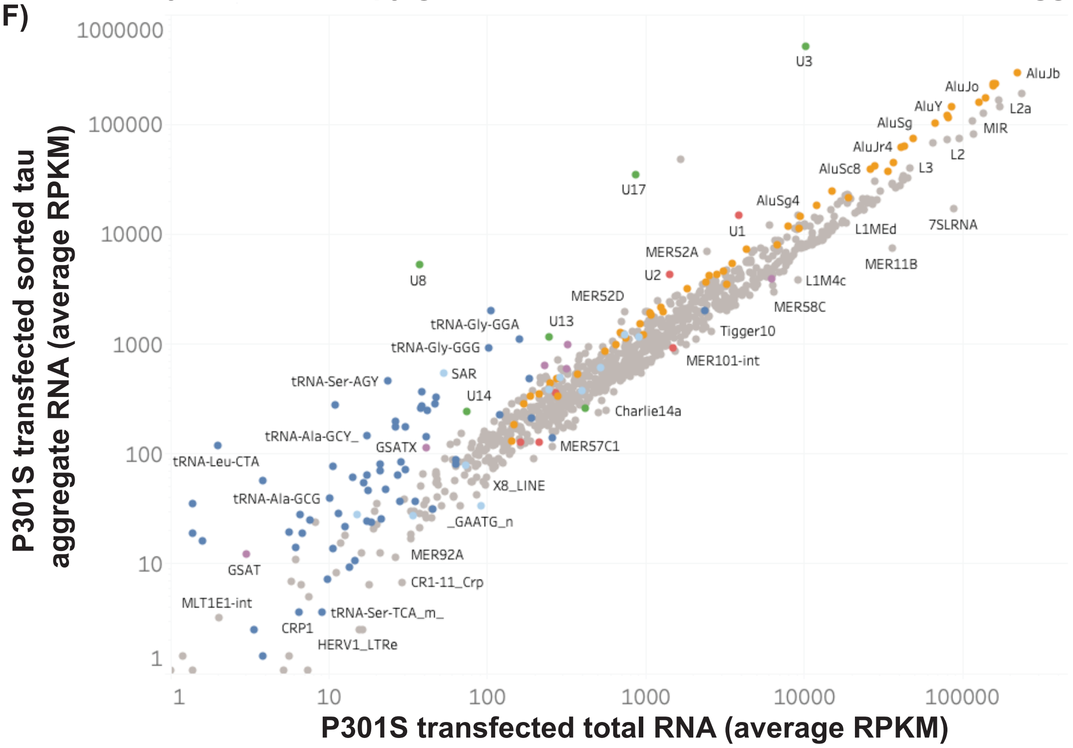


G)

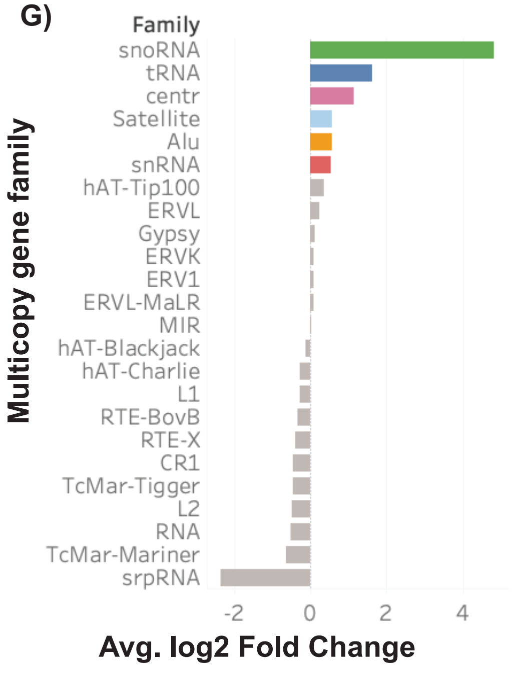

### Supplemental Figure 2 continued: Additional data on HEK293 and mouse tau aggregate isolation and sequencing

H)

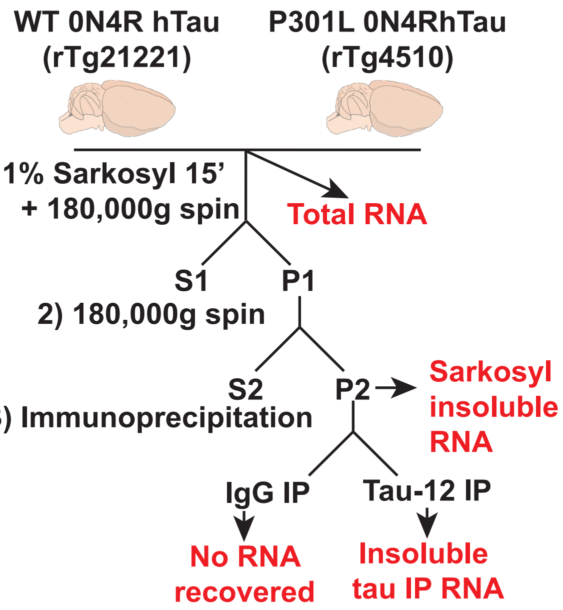


I)

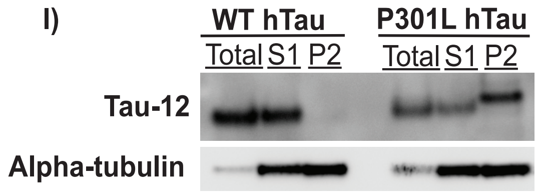


J) P301L vs WT sarkosyl insoluble gene type enrichment

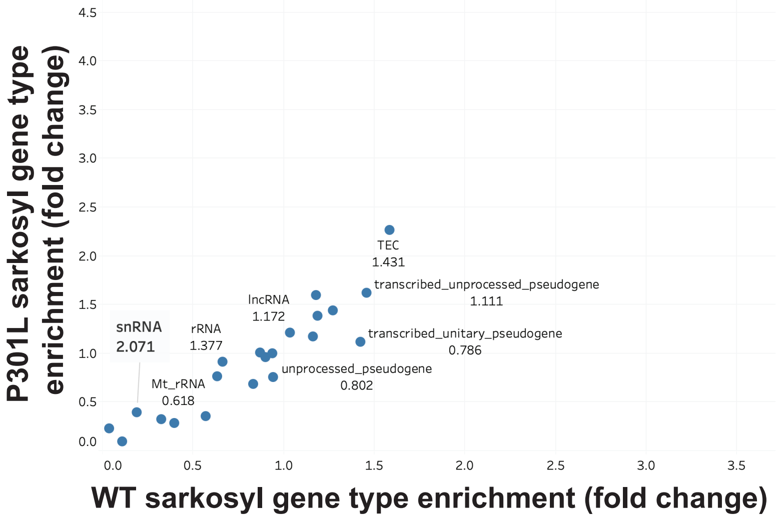


K) Mouse vs HEK293 gene type enrichment

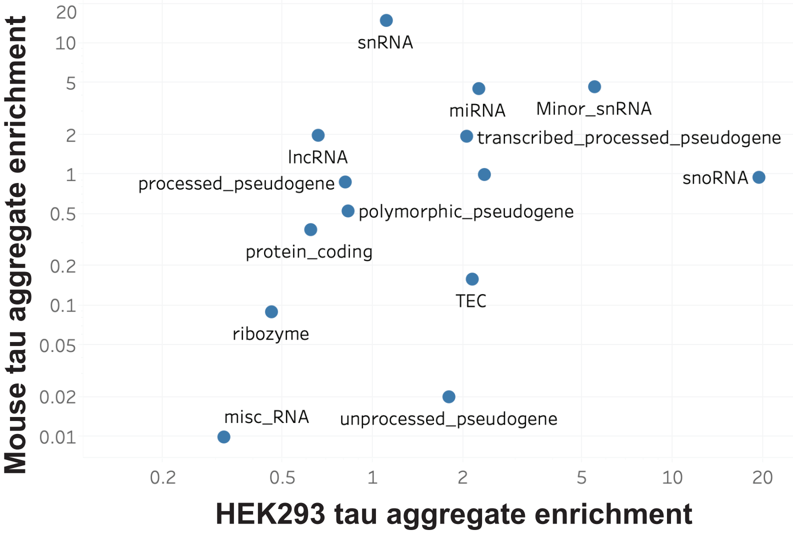


L) mRNAs significantly enriched in tau aggregates isolated from mouse brains

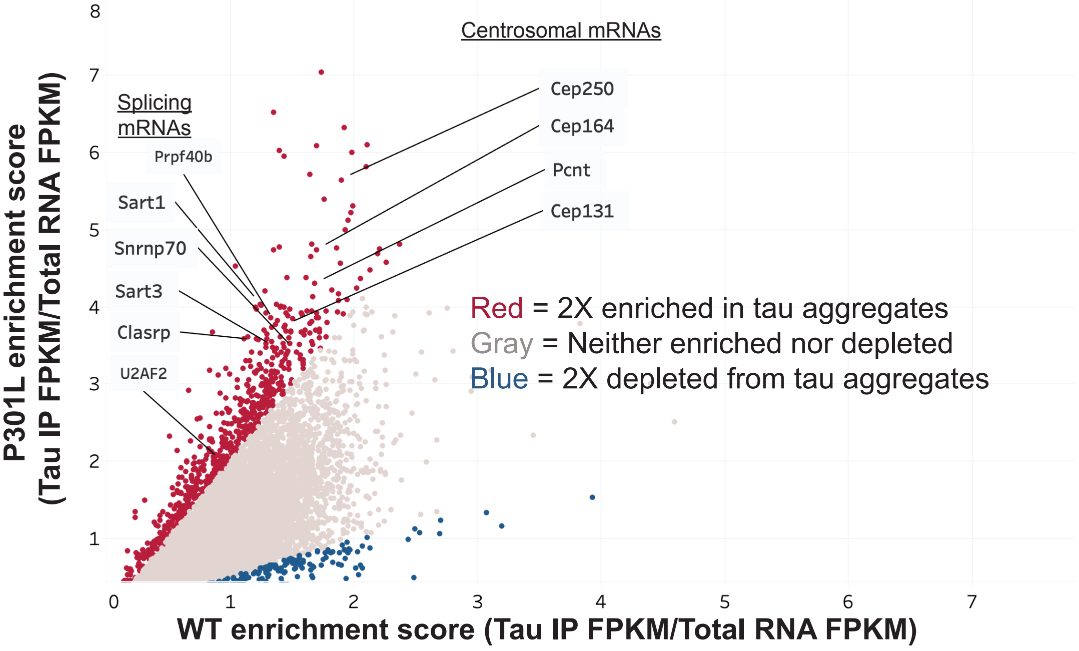

**Supplemental Figure 3: Additional FISH for RNAs in HEK293 tau biosensor cells**

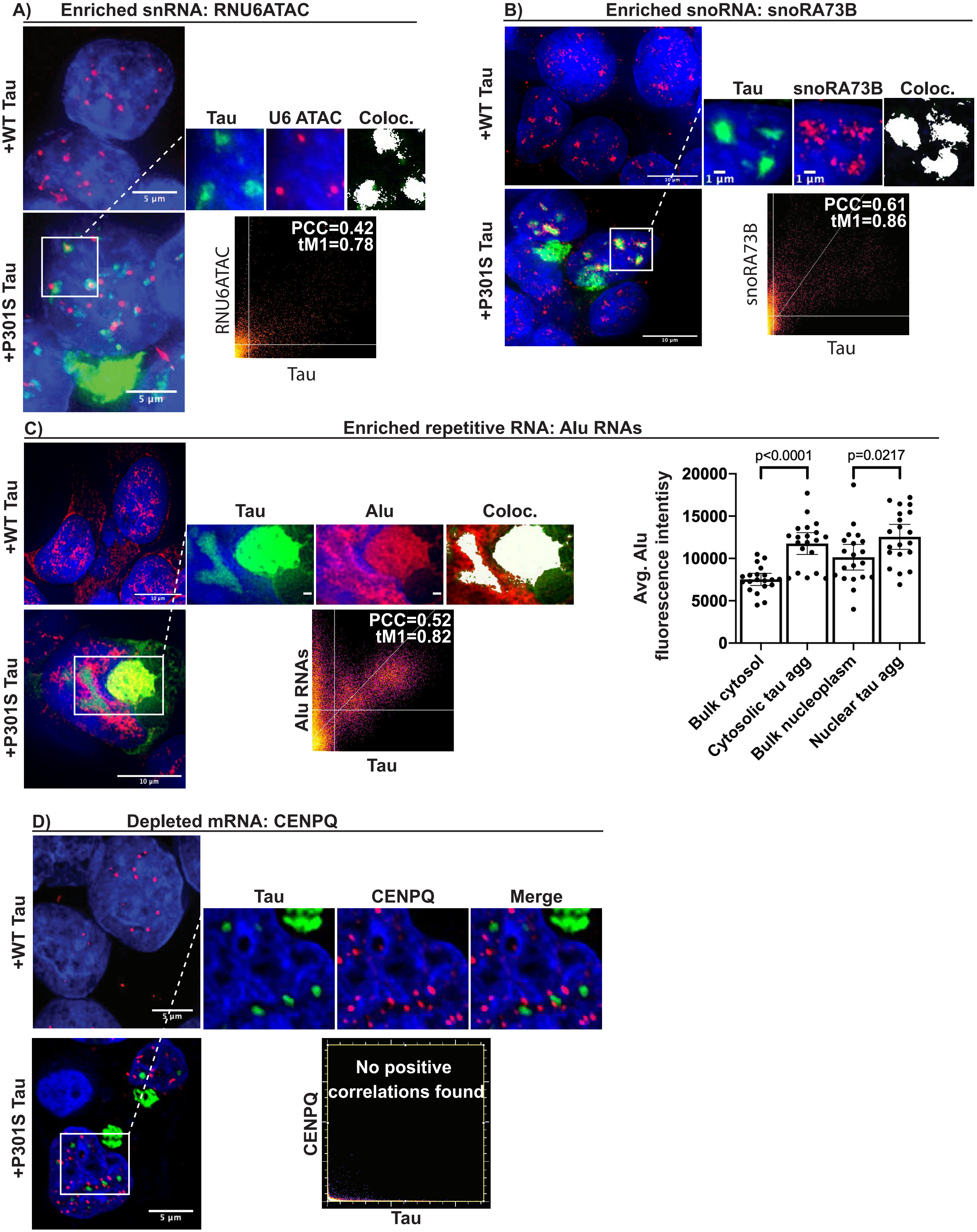

Supplemental Figure 4 : Additional HEK293 tau biosensor cell data

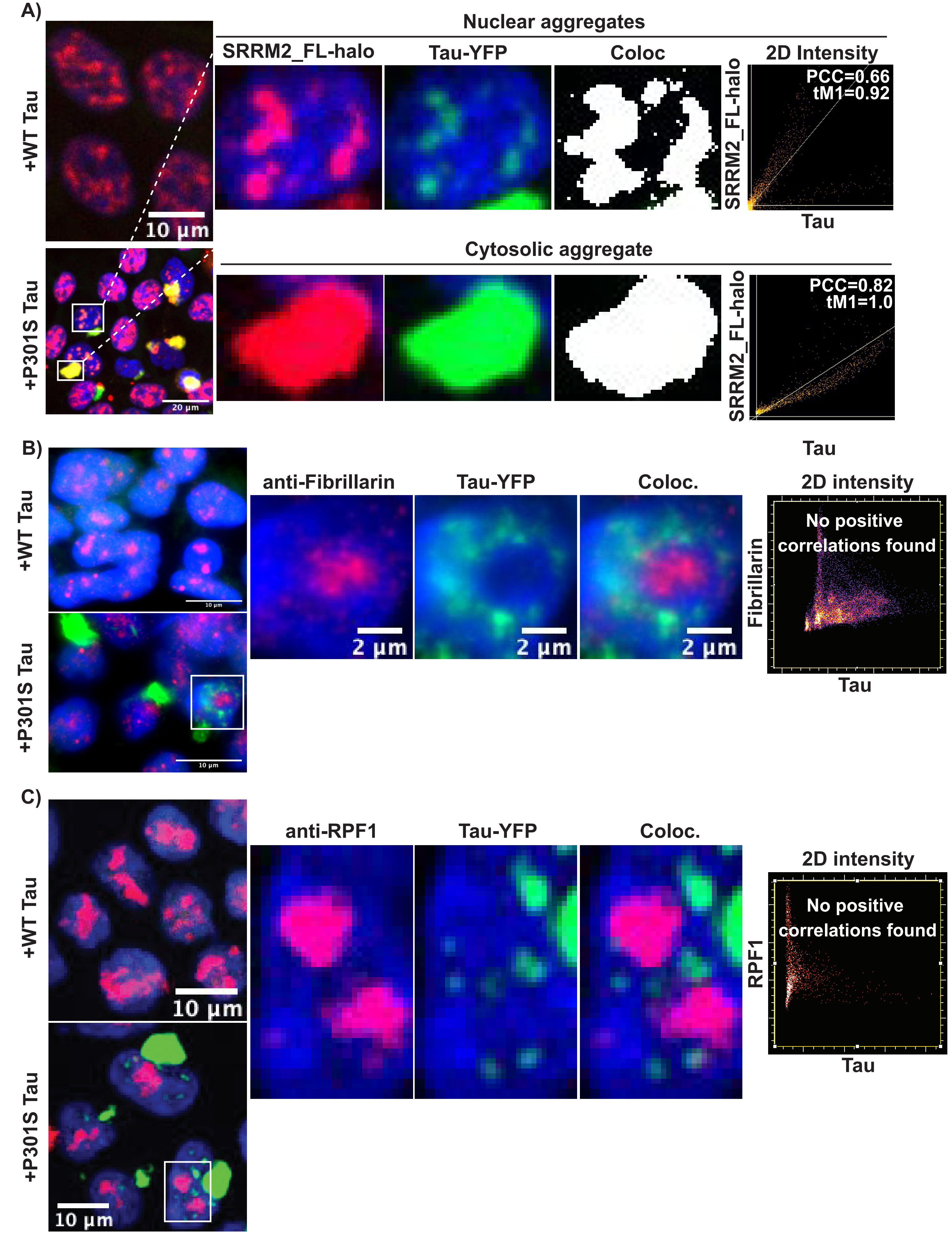

Supplemental Figure 4 continued: Additional HEK293 tau biosensor cell data

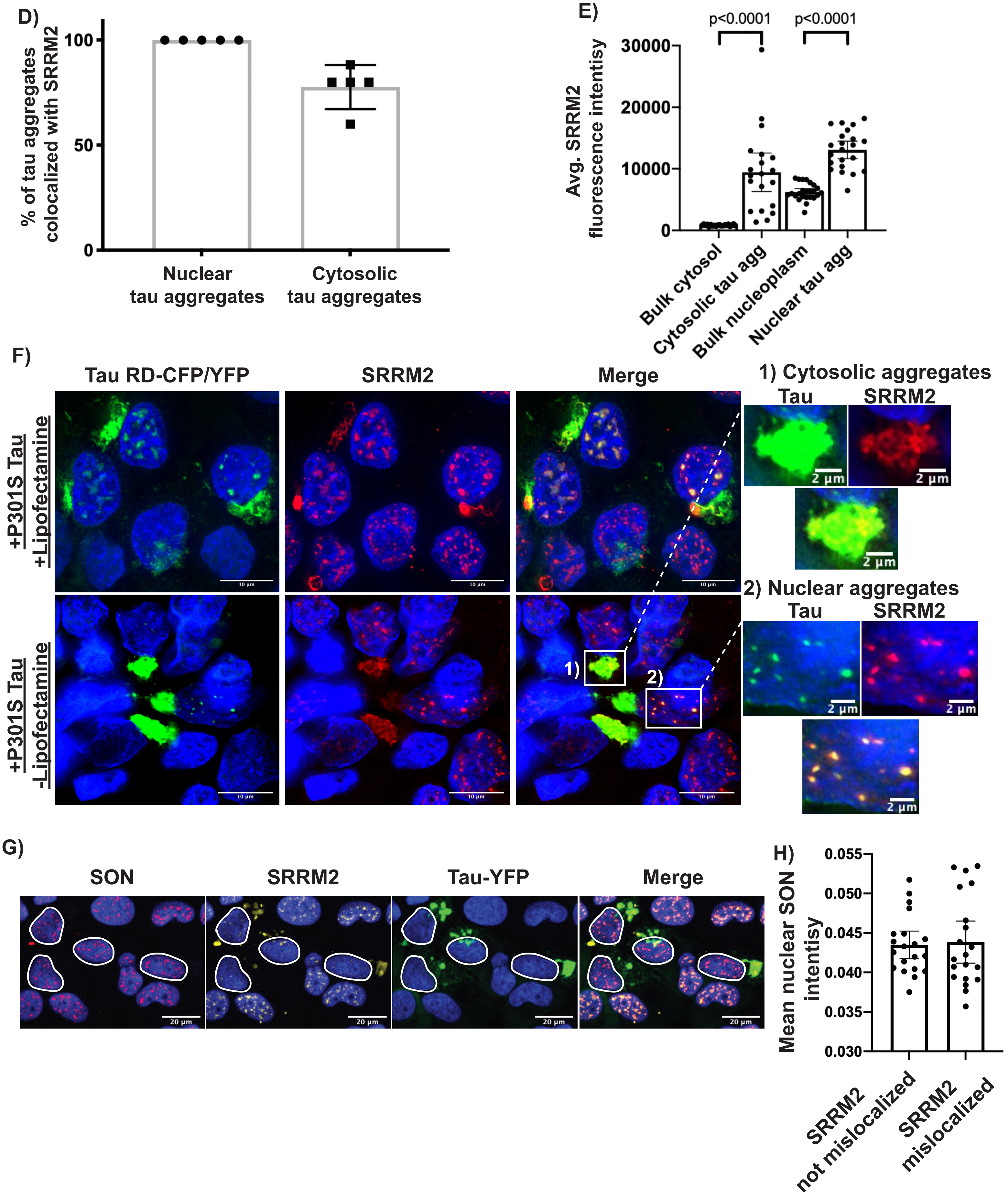

Supplemental Figure 5

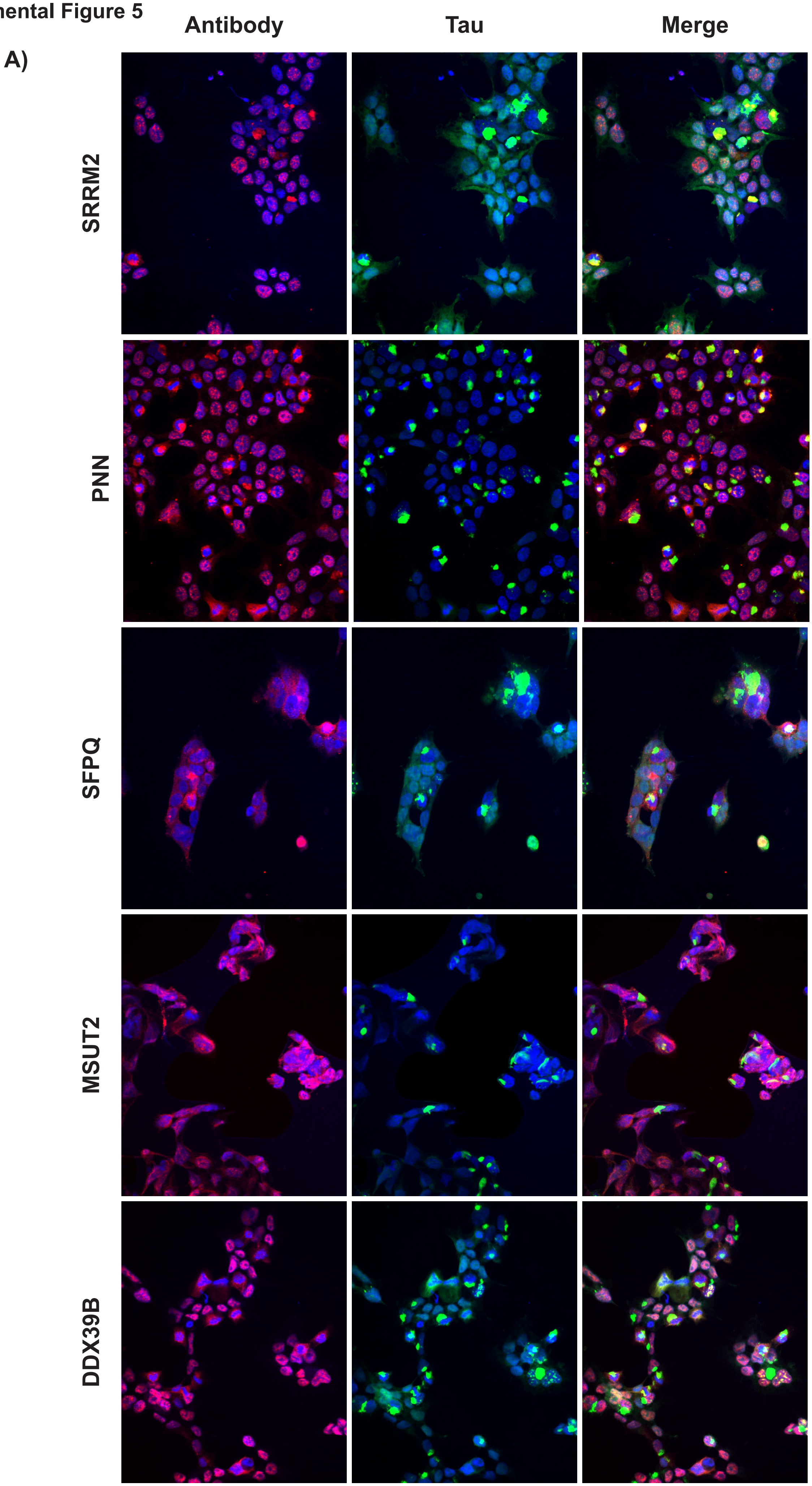

Supplemental Figure 5 continued

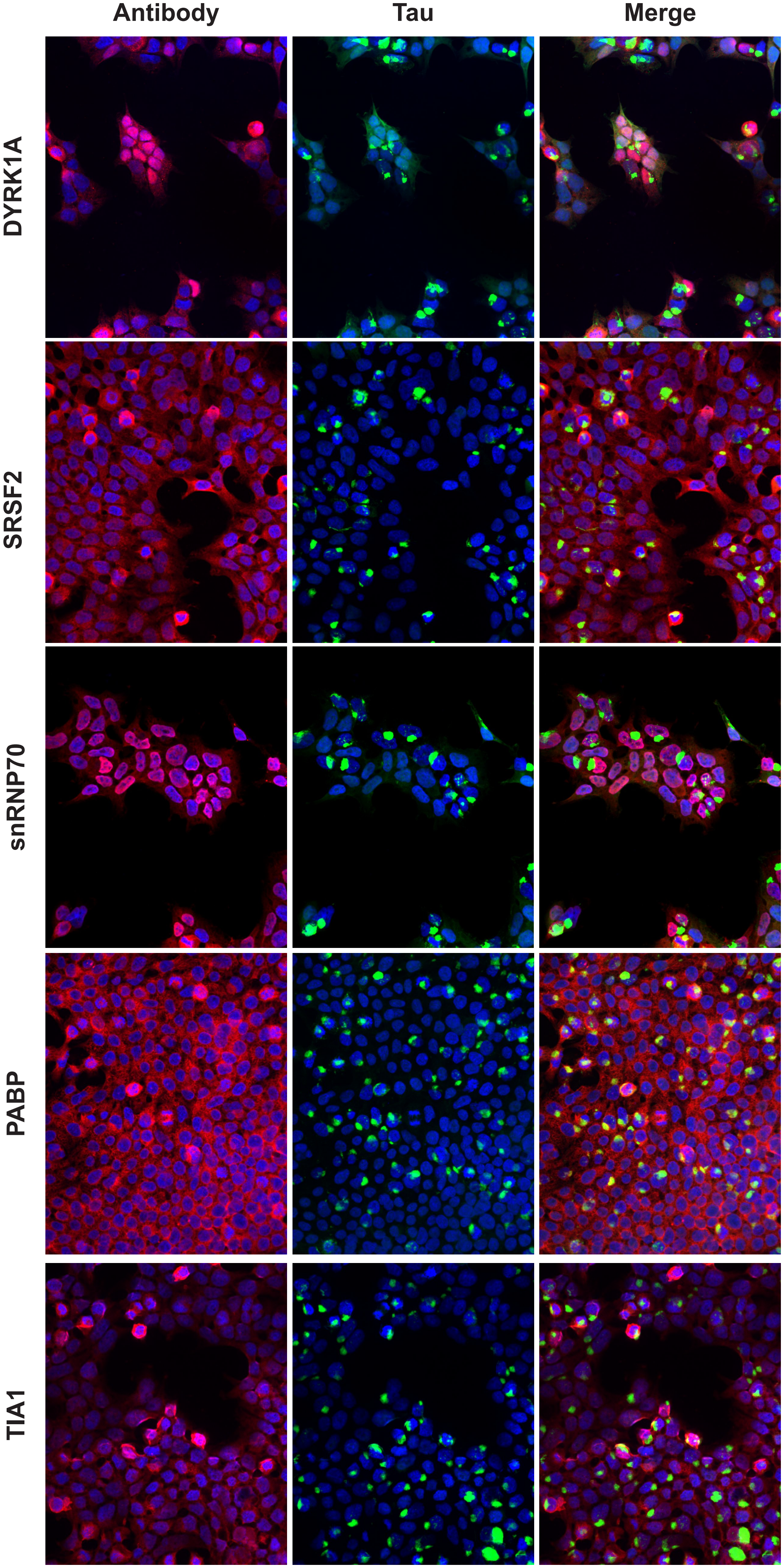

Supplemental Figure 5 continued

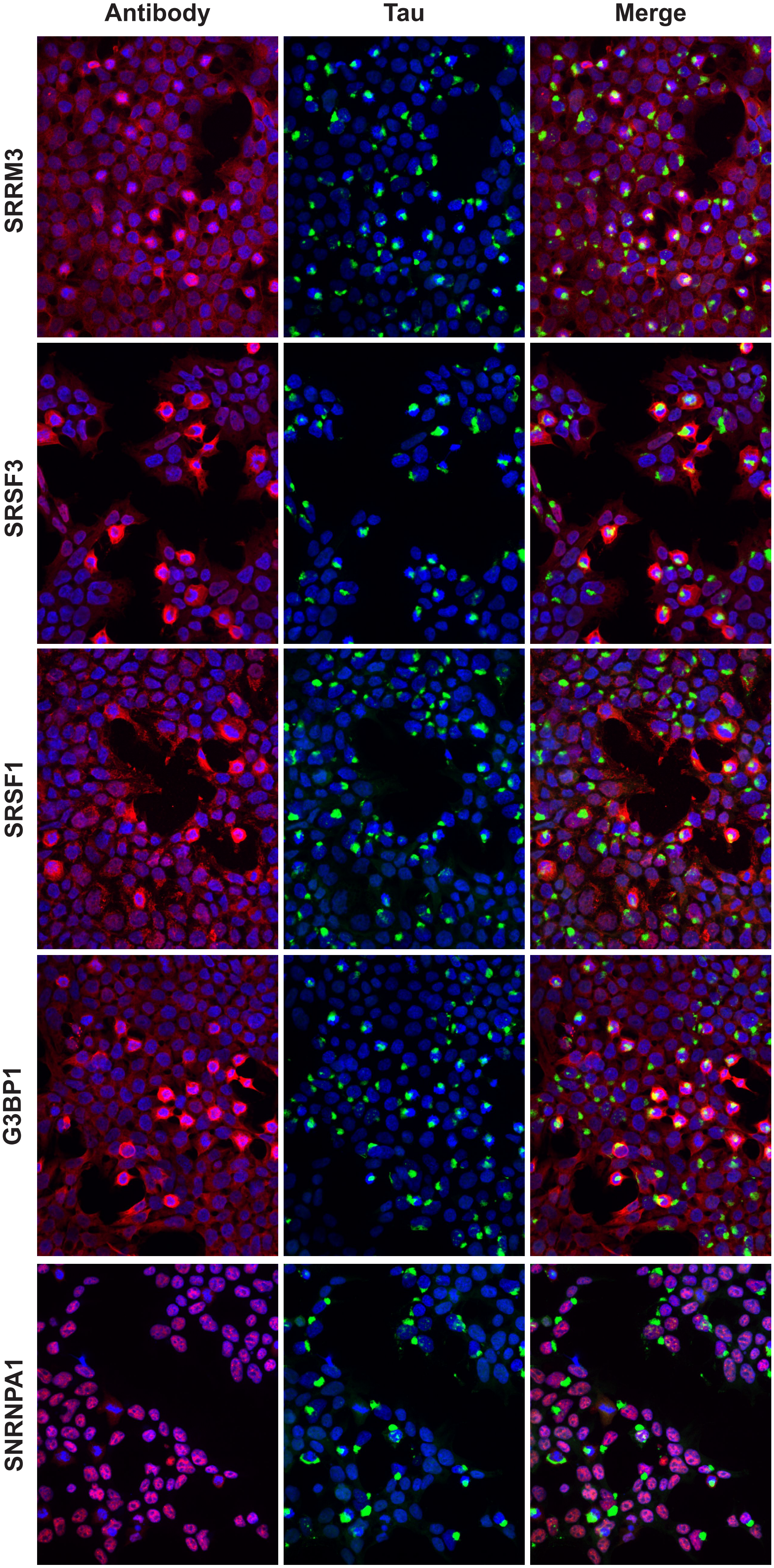

Supplemental Figure 5 continued

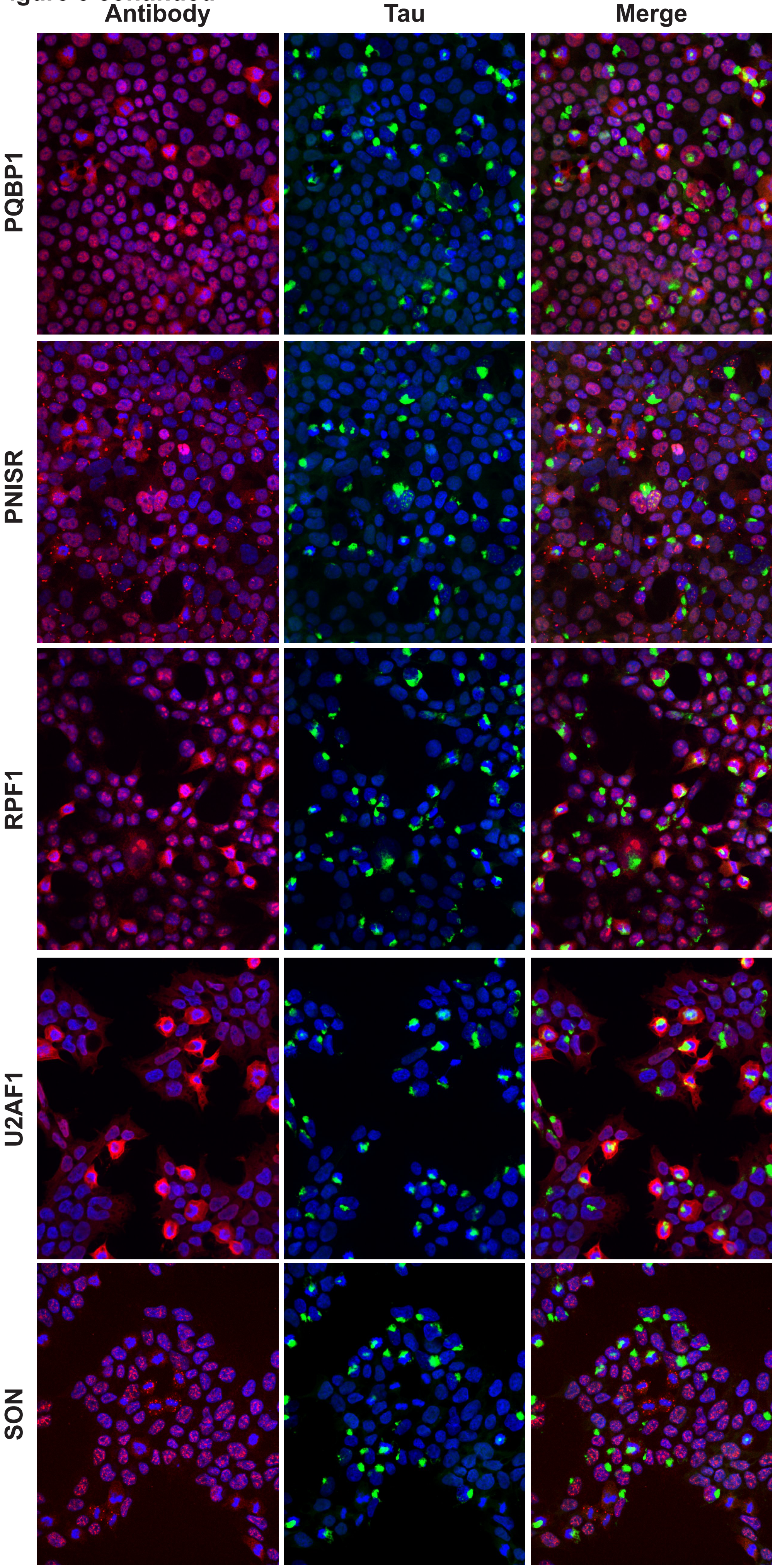

**Supplemental Figure 5 continued**

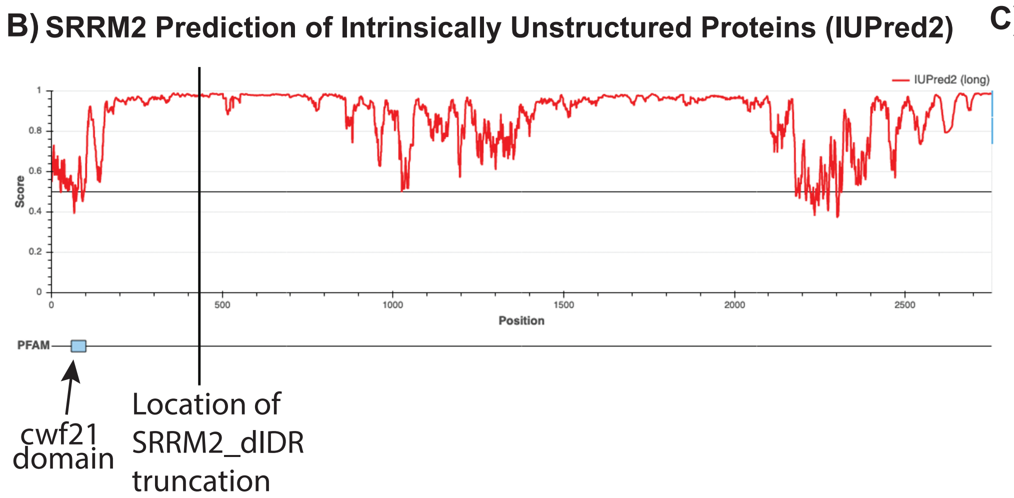


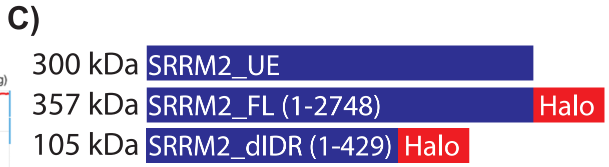


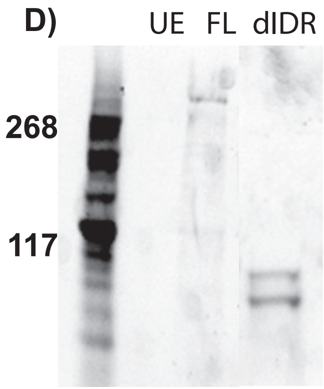

**Supplemental Figure 6: Additional splicing analysis data from cells with and without tau aggregates**

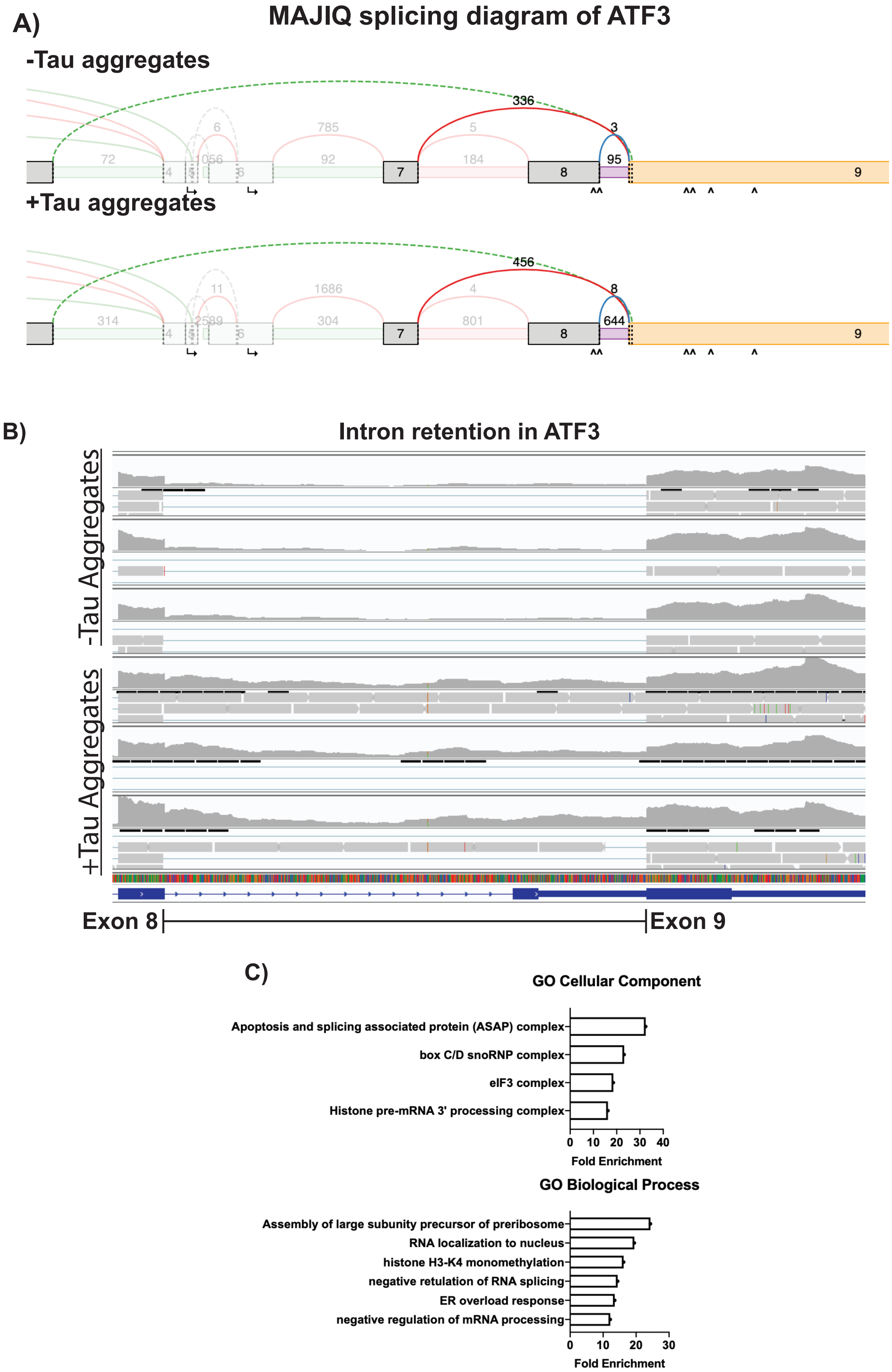

**Supplemental Figure 7: Additional in vivo data from mouse and human tauopathies**

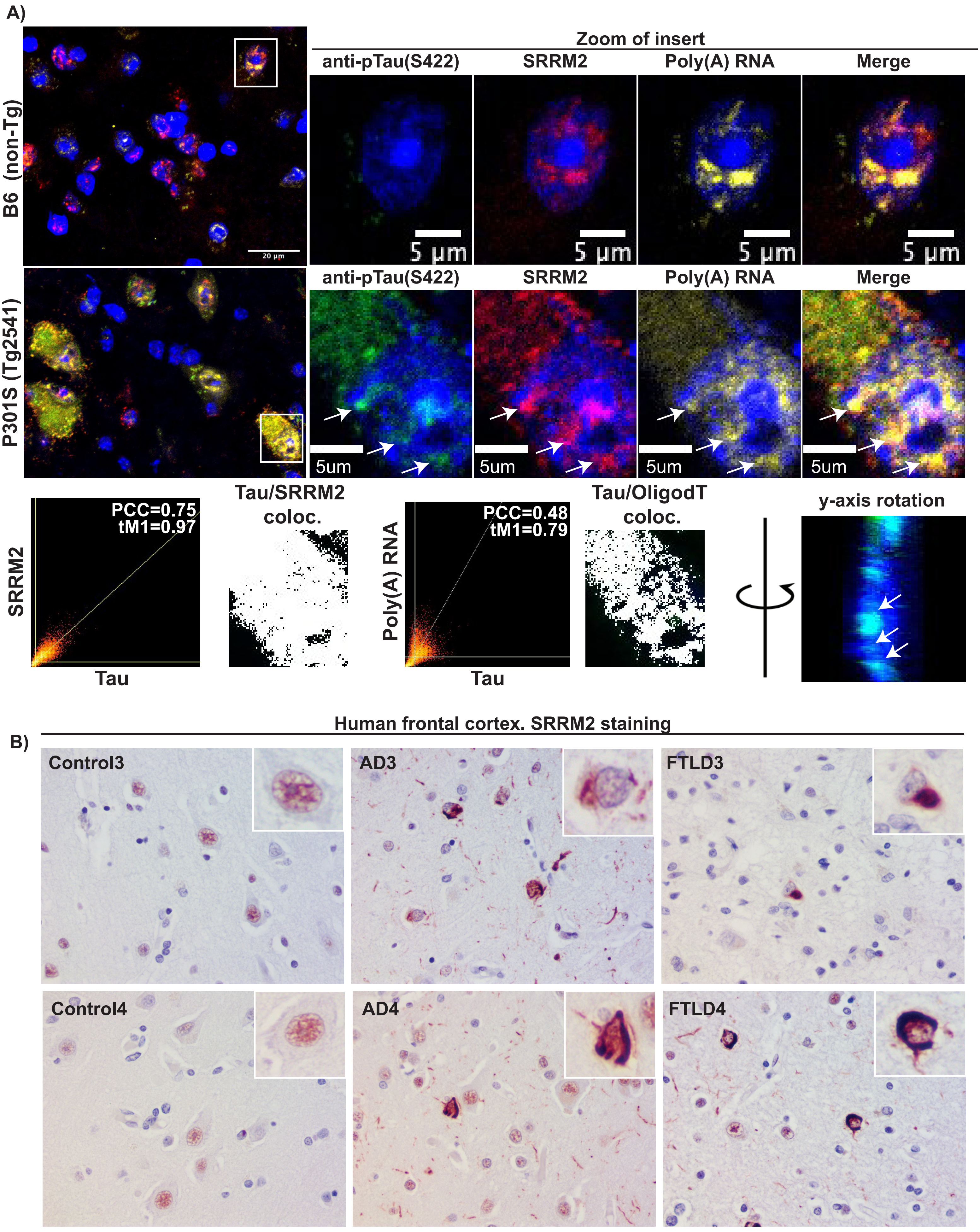
